# *k*ache-hash: A Dynamic, Concurrent, and Cache-Efficient Hash Table for Streaming *k*-mer Operations

**DOI:** 10.64898/2026.02.13.705625

**Authors:** Jamshed Khan, Rob Patro, Prashant Pandey

## Abstract

Many computational genomic analyses spend much of their time inserting and looking up *k*-mers in large hash tables. These operations stream through sequences, and consecutive *k*-mers share k − 1 characters. General hash tables ignore this locality. Each *k*-mer costs them a memory access, and in concurrent builds, a synchronization. We present *k*ache-hash-hash, an exact, dynamic, concurrent, and resizable hash table that places *k*-mers by their minimizers.

A run of consecutive *k*-mers with the same minimizer then shares its buckets, which it fetches and locks only once. Building on Iceberg hashing, *k*ache-hash handles skewed minimizers with secondary buckets. A compact spill filter lets lookups stop early, yet stay exact. The table stores sets or maps, and doubles in place in parallel. Under a cache-residency assumption, we show that a run that stays in its primary bucket brings in at most that bucket’s cache lines, however long the run is.

We compare *k*ache-hash with six general-purpose and *k*-mer-specific tables on five genomic datasets. At 56 threads, it builds *k*-mer sets 3.5–7.0× faster than the fastest baseline, and answers queries 1.5–5.4× faster. On the human genome, its positive and mixed queries fetch 3.1–3.2× fewer cache lines per *k*-mer than those of the fastest baseline. In PanGenie, *k* ache-hash speeds up end-to-end genotyping 1.96× with identical outputs. In `K2Rmini`, it speeds up read filtering 10.7–19.4× over a fixed control version of the release, also making the filter exact. *k*ache-hash is implemented in C++20 and is available at https://github.com/jamshed/kache-hash.

## 1 Introduction

Hash tables of *k*-mers, the substrings of length k, are at the core of many genomic analyses and can dominate their runtime. They underpin *k*-mer counting [30,37], read alignment [25,3], variant genotyping [17,10], RNA quantification [38], metagenomic classification [50], and sequence search [31]. `PanGenie` genotypes a sample by counting the *k*-mers of a pangenome in its reads [10]. This counting takes 54% of its genotyping time in our experiments (§5.1). `K2Rmini` filters reads by the share of their *k*-mers found in a reference [34]. Building its *k*-mer index takes over 94% of its runtime (§5.2). Such tables must thus be fast to build and query.

These workloads stream through sequences, one *k*-mer after the next. Consecutive *k*-mers overlap in k − 1 characters, and often share their *minimizer*, the ℓ-mer with the least hash [42,43]. A general hash table ignores this, placing each *k*-mer by its own hash. Once the table outgrows the caches, nearly every *k*-mer then brings in a cache line of its own from memory. A concurrent build also synchronizes on every *k*-mer. If *k*-mers were placed by their minimizers instead, a *run* of consecutive *k*-mers with the same minimizer would share a bucket, and with it the memory traffic and the locking.

Existing *k*-mer structures exploit this locality in static indexes, or to save space. `SSHash` groups *k*-mers by minimizer for compact storage and fast streaming queries, but is static [39]. `SBWT` [2], `FMSI` [45], and `LPHash` [40] exploit the overlaps of *k*-mers to save space. `CBL` [32] and `Brisk` [46] use it to keep dynamic *k*-mer sets compact. Fast concurrent tables, from `Jellyfish`’s counting table [30] to general-purpose ones such as `IcebergHT` [36], `libcuckoo` [26], and `Boost` [7], place each *k*-mer by its own hash. `K2Rmini`’s authors name its *k*-mer set as its main limitation, and propose a concurrent table that groups *k*-mers by minimizer in its place [34].

We present *k*ache-hash, an exact, dynamic, concurrent, and resizable hash table for *k*-mer sets and maps, designed for throughput (§2). It builds on Iceberg hashing [5], whose stable placement and few candidate buckets limit data movement and probes [36]. Unlike Iceberg hashing, it picks a *k*-mer’s buckets by its minimizer. The run, not the *k*-mer, thus becomes the unit of work. Its buckets are fetched once, and an insertion locks them once.

But grouping keys breaks the independence that hash tables rely on. Minimizer frequencies are skewed [29], so buckets fill unevenly. When a *k*-mer’s primary bucket is full, the *k*-mer goes to the emptier of two secondary buckets in the same table, or else to a small overflow table. Lookups of absent *k*-mers would then have to probe further. Instead, a small *spill filter* in each bucket marks the minimizers that spilled. A lookup can then stop at the primary bucket, and still be exact. The table also doubles in place, in parallel, and moves most keys without locks.

As keys no longer hash independently, we bound costs per run (§3). We count the cache lines that operations bring in, prefetches included. A run that stays in its primary bucket then brings in at most that bucket’s lines, five in a set, however long the run is. In a random sequence, this is at most about 0.91 lines per *k*-mer, for k = 31 and ℓ = 22. Runs that spill cost more, but still *O* (1) expected lines per *k*-mer.

We compare *k*ache-hash with six other tables on five datasets, from a human genome and human reads to collections of *E. coli* and gut genomes (§4). At 56 threads, it builds *k*-mer sets 3.5–7.0× faster than the fastest baseline on each dataset, presized or resizing, and answers queries 1.5–5.4× faster. Its queries even outperform `SSHash’s` static index, by 2.2–8.0×. It leads on one thread too, by 1.8–4.9 ×. From 1 to 56 threads, its presized builds speed up 34–37×, and its queries 29–43×. On the human genome, its builds and positive and mixed queries fetch 0.84–1.02 cache lines per *k*-mer, spills included, against 1.5–5.1 for the other dynamic, concurrent tables. These run at 77–85% of the memory’s bandwidth for random access. Presized, its sets take 13.0–13.3 bytes per distinct *k*-mer, less than the other dynamic, concurrent tables, except `Jellyfish` on three datasets.

These gains carry over to real tools (§5). In `PanGenie` and `K2Rmini`, we swap *k*ache-hash in for the *k*-mer table, and parallelize the steps around it that would become the bottleneck. `PanGenie` then genotypes a sample 1.96 × faster, with identical outputs. Against a control with the same parallel steps, the table alone gives 1.50 ×. `PanGenie’s` CPU time per sample also drops from 12.3 to 4.2 hours, saving about 26,000 CPU-hours over its 1kGP callset. `K2Rmini` runs three filtering tasks 10.7–19.4× faster than its release with its minimizer set’s skewed hash fixed. It also becomes exact, without the false matches of its 32-bit *k*-mer hashes.

## 2 *k*ache-hash: Design and Implementation

*k*ache-hash groups the operations on consecutive *k*-mers that share a minimizer, to reuse cache lines and amortize locking. It is a dynamic, concurrent and resizable hash table of *k*-mers. It stores a set of *k*-mers, or a map from *k*-mers to values. Its operations take whole sequences, and go through their *k*-mers in order.

*k*ache-hash builds on Iceberg hashing [5,36]. Iceberg hashing hashes a key to one bucket of a *front yard* of large buckets. If that bucket is full, the key goes to a small *backyard*: to the emptier of two more buckets, or if both are full, to an overflow table. Keys stay in place until the table resizes, and each key has only a few possible placements. *k*ache-hash follows a similar but merged leveling structure, and picks a *k*-mer’s buckets by its minimizer. Consecutive *k*-mers often share their minimizer, and thus their buckets. A run of such *k*-mers, not a single *k*-mer, becomes the unit of work. Its buckets enter the cache once, are locked once, and are prefetched once (Fig. 1).

**Fig. 1.**
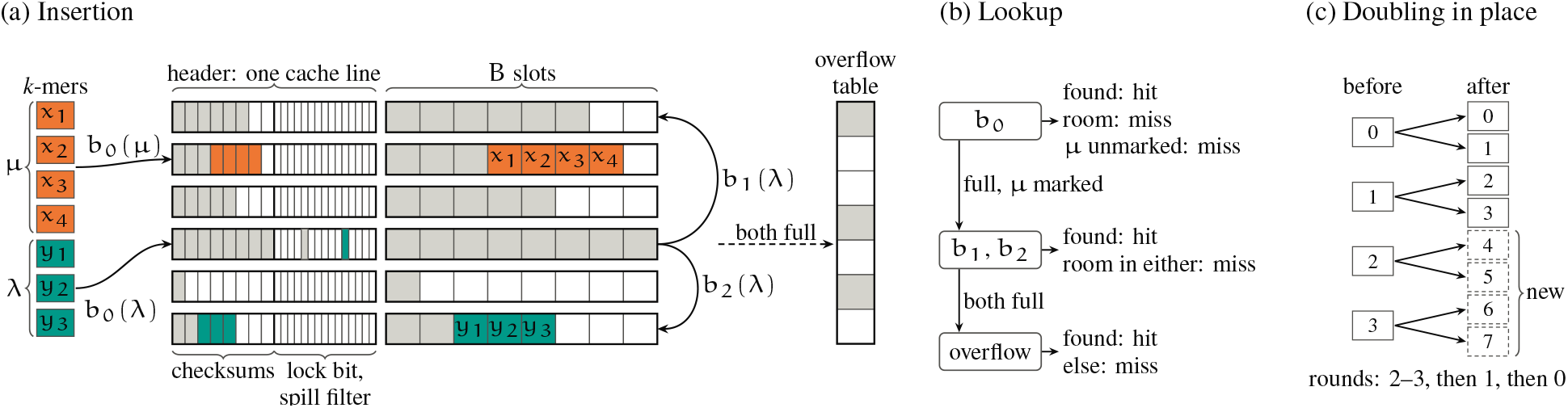
*k*ache-hash’s table, drawn with B = 8 slots per bucket instead of 32. A bucket is a row, containing a header of checksums and flags, then its slots. Gray slots hold existing keys, and white ones are empty. (a) Inserting two runs of *k*-mers, of the minimizers µ and λ. The run of µ fills free slots of µ’s primary bucket, b_0_(µ). The primary bucket of λ, b_0_(λ), is full. Hence λ is marked in the bucket’s spill filter, and the run goes to the emptier secondary bucket, b_2_(λ). (b) A lookup’s probes, and its exits. (c) Doubling in place: the keys in their primary bucket b move to 2b or 2b + 1, in rounds from the top.

### 2.1 Table Structure

*k* ache-hash has a *main table* of m buckets and a small *overflow table* (Fig. 1a). A *k*-mer has a *primary* bucket and two *secondary* buckets in the main table. Unlike Iceberg hashing’s backyard, the secondary buckets are buckets of the main table, so a bucket holds keys of both kinds.

#### *k*-mer encoding

*k*ache-hash-hash’s design and implementation restrict k ≤ 32. With 2-bit encoding of characters, a *k*-mer fits in a 64-bit machine word. A cache line can thus hold 8 *k*-mers.

#### Buckets

A bucket has B = 32 slots. Each slot holds a key, the *k*-mer, and in a map, also its value. A bucket starts with a 64-byte *header*, aligned to a cache line. Its slots then follow. A set’s bucket thus spans five cache lines. The header holds a one-byte *checksum* for each slot, then 256 flag bits. A checksum is an 8-bit hash of the slot’s key, from 1 to 255, and 0 marks an empty slot. Flag 0 is the bucket’s lock (§2.4), and the other 255 flags form its *spill filter* (§2.3).

#### Probes

To search a bucket for a key, *k*ache-hash compares its 32 checksums with the key’s checksum in one 256-bit vector instruction. It then compares the key only with the keys matching the checksum. For an absent key, a bucket of s keys has s/ 255 < 0.13 false matches in expectation. A checksum comparison against 0 finds the first empty slot. *k*ache-hash deletes no keys, so a bucket’s slots are filled in order. A bucket is thus full exactly when its last checksum is non-zero. A probe usually touches the header’s cache line, and at most one line of slots.

#### Capacity and overflow table

A table presized for n keys at *load factor* α is sized to the fewest buckets m with αmB > n. The overflow table is a concurrent linear-probing table [22], with a lock per slot. It stores its keys complemented, so zeroed memory holds empty slots.^3^ It starts with room for 1 /32 as many keys as the main table. It doubles on its own once it reaches its load factor, so skew does not make the main table resize early.

### 2.2 Minimizer-Based Placement

A *k*-mer and its reverse complement are one key, stored as the smaller of their encodings: the *canonical* form. The *minimizer* of a *k*-mer is its ℓ-mer whose canonical form has the least hash [42,43]. *k*ache-hash’s ℓ-mer hash function is a bijection, so the least hash h identifies the minimizer µ. A *k*-mer and its reverse complement share µ, so a key’s placement does not depend on its strand.

A *k*-mer has w = k − ℓ + 1 many ℓ-mers. Consecutive *k*-mers share w − 1 of them, so the *k*-mers often share their minimizer. We call a maximal stretch of consecutive *k*-mers with one minimizer a *run*, also known as a super-*k*-mer [27].

*k*ache-hash derives a *k*-mer’s buckets from its minimizer’s hash h. The primary bucket is

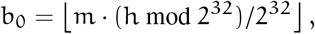

a multiply-and-shift range reduction that works for any m [24]. It uses the low bits of h, as the least of w hashes has high bits that lean toward 0. Doubling m takes b_0_ to 2b_0_ or 2b_0_ + 1, which matters in resizing (§2.5). The secondary buckets, b_1_ and b_2_, come from two more hashes of h. The *k*-mer itself determines only its checksum and its slot in the overflow table. All the *k*-mers of a run thus have the same buckets. The run’s first *k*-mer brings them into the cache, and the others find them there. §3 estimates the runs’ lengths, and bounds the resulting cache misses. **Skew**. Minimizer frequencies are skewed [29], so this placement loads the buckets unevenly. A frequent minimizer may correspond to more *k*-mers than a bucket holds. Its primary bucket then fills, and its other *k*-mers go to its secondary buckets or the overflow table. Even then, the run’s *k*-mers share their secondary buckets.

### 2.3 Streaming Operations

An operation splits its sequences into *stretches* at characters other than A, C, G and T.

#### Insertion

To insert a *k*-mer x with minimizer µ, *k*achehash first probes b_0_ (Fig. 1a). If x is there, a set’s insertion ends, and a map’s applies the caller’s update function to the value. Otherwise, if b_0_ is not full, *k*ache-hash writes x there, and then its checksum. If b_0_ is full, *k*ache-hash marks µ in b_0_’s spill filter, at a flag picked by other bits of h. It then probes b_1_ and b_2_. If x is in neither, it goes to the emptier one, by the power of two choices [4,41]. If both are full, x goes to the overflow table.

#### Lookup

A lookup takes the same path, but can stop early (Fig. 1b). An insertion goes past a bucket only if the bucket is full, and buckets only gain keys. So if b_0_ has an empty slot now, it had one whenever a *k*-mer of µ was inserted. If present, x must then be in b_0_. Likewise, if b_0_ is full but b_1_ or b_2_ has room, x is not in the overflow table. The spill filter adds a third exit. An insertion marks µ in the filter before it places a *k*-mer of µ past b_0_. So if b_0_ is full but does not mark µ, x is absent. This exit matters for absent *k*-mers, as skew leaves many primary buckets full. Without it, each such lookup would probe two more buckets, and often the overflow table. With the filter, these probes are incurred for an absent minimizer only for false marks from some other minimizer. It acts as a 255-bit single hash Bloom filter [6]. Its false marks cost only probes, never wrong answers, as lookups compare full keys. Appendix B.1 proves the exits correct.

### 2.4 Concurrency

Insertions lock the buckets that they probe, and lookups take no locks.

#### Bucket locks

A thread takes a bucket’s lock, flag 0 of its header, with an atomic compare-and-swap. A waiting thread spins on plain loads, and retries only once the lock looks free [14]. An insertion locks a bucket even to search it, as otherwise two threads inserting the same new *k*-mer could both find it absent, and both place it.

#### One lock per run

An insertion locks a run’s primary bucket at the run’s first *k*-mer, and keeps it until the run ends. The lock, like the cache misses, is thus paid once for a run that stays in this bucket. If the bucket is full, the thread releases it, and locks all three of the run’s buckets for the rest of the run. The full primary bucket is locked again since the run’s later *k*-mers may be in it, and a map updates their values there. Locking is done in increasing order of index, which rules out deadlocks. The thread releases the buckets before it goes to the overflow table, which locks its own slots. It locks the three buckets again for the run’s next *k*-mer.

#### Lookups without locks

An insertion writes a key before writing its checksum, and marks a spill before it places a key past the primary bucket. So a concurrent lookup either finds a key, or misses it as if the lookup came first, linearizing before the insertion [14]. A map’s inplace update inserts nothing, and takes no locks either. It changes its value with an atomic compare-and-swap loop.

#### Counting keys

To track the size of the table in distinct keys, each thread counts its new keys privately, and adds them to the shared count after at most 2^14^ of them.

### 2.5 Resizing in Place

*k*ache-hash doubles its main table once its count of keys reaches αmB (§2.1). The thread that finds this threshold crossed, while counting its inserted keys, executes the resize with a pool of workers.

#### Table lock

Insertions hold a *readers–writer lock* [14] in shared mode (reader), and a resize holds it exclusively (writer). An insertion thread marks its read-lock intent in its own flag, on its exclusively owned cache line, rather than in a shared count. An insertion yields to a waiting resize at each run’s start, when it holds no bucket locks. **Growing in place**. In resizing, the main table’s space is doubled in place.^4^ The new half reads as zeros, which are empty buckets. A resize thus never holds two main tables separately.

#### Moving keys

Resize completes in two phases. The first phase moves the keys that are in their primary buckets, and the second moves the rest. The first phase moves each key in its primary bucket b to its new primary bucket, 2b or 2b + 1 (Fig. 1c). No other old bucket maps to them, so the workers take no locks. The old buckets go in rounds, from the top buckets to the bottom, so that no key lands in a bucket whose keys are yet to move. Starting from e = m, a round moves the buckets [s, e) into [2s, 2e), where s = ⌈e/ 2⌉, and the next round starts from e = s. The buckets [2s, 2e) are either new or emptied in earlier rounds. Once e = 1, one worker splits bucket 0 into buckets 0 and 1, compacting the keys that stay. This phase also clears the spill filters, and sets aside the keys that were in their secondary buckets in a small buffer of its own. In the second phase, the workers insert these keys and the overflow table’s keys, with locks, into the doubled table and a new overflow table. Many of them now fit in their primary buckets, and the others mark the filters. The new overflow table is sized afresh to either 1 /32 as many keys as the doubled main table, or as many as the old one, whichever is more.

#### Recomputing minimizers

A bucket’s keys do not form a sequence, so their minimizers cannot be rolled along one. A resize instead recomputes each key’s minimizer hash, four keys at a time in SIMD lanes. Storing each key’s minimizer position would save this work, but would either take the header’s room for the spill filter, or add to the header’s size.

### 2.6 Implementation

#### Interface

*k*ache-hash is implemented as a header-only C++20 library. Template parameters fix k, ℓ, the value type (none for a set), and whether the table is concurrent. A C interface with Rust bindings (§5.2) and a counting library (§5.1) serve other applications. Appendix <u>A</u> lists the operations and their concurrency rules, and adds details on resizing.

#### Describing, then operating

An operation first *describes* the *k*-mers of its input in chunks: their canonical forms, checksums and minimizer hashes. It describes each chunk ahead of operating on it. Describing many *k*-mers at once vectorizes well, and informs the table of the coming operations’ tentative probes.

#### Minimizer computation

The describer rolls the 2-bit forms of the *k*-mer and of its last ℓ-mer, with their reverse complements, along a stretch. It hashes each canonical ℓ-mer with Wang’s 64-bit integer hash [47], which uses only shifts, additions and exclusive-ors. It computes the minimum of a *k*-mer’s window of w hashes by doubling. ⌊log_2_ ⌋w rounds of pairwise minima compute the minima of spans with powers-of-two lengths. Two such spans cover each window. This way, all the minimizer computations across a stretch of *k*-mers are independent, branch-free, and vectorized. *k*ache-hash describes four stretches at once with AVX-512, one per 64-bit lane of 256-bit vectors. Long stretches split four ways; short ones, such as reads, go four at a time. Vectors stay at 256 bits, without 64-bit multiplies, which keeps the cores at their full clock speed.

#### Lookahead prefetching

*k*ache-hash prefetches in two stages. Once a chunk is described, it prefetches the headers of the chunk’s primary buckets. Ahead of an operation, it reads the header, and prefetches the slot that the operation will likely touch. That is the slot whose checksum matches, or for an insertion, the first empty one. If no check-sum matches and the bucket is full, an insertion instead prefetches ahead the secondary buckets’ headers, once per run, and the *k*-mer’s overflow slot. Lookups prefetch less. They suppress slot prefetches after misses, and prefetch secondary headers only if the spill filter marks the minimizer. The beginning of a run spares its later *k*-mers their cache misses, and the prefetches hide the latency of following runs’ memory traffic.

#### Memory

*k*ache-hash maps its tables on 2 MB transparent huge pages to reduce translation costs of random addresses, and prefaults each page in parallel during table allocation. `mremap` keeps the huge pages through resizes. Zeroed memory holds empty slots in the main and the overflow tables, so neither needs an initialization pass. Map slots pack keys and values bytewise without padding: a one-byte value adds one byte to a slot, rather than eight in an ordinarily aligned key–value pair.

## 3 Guarantees and Costs

This section bounds the cache lines that operations bring in. Iceberg hashing bounds how many keys leave its front yard, as its keys hash independently [5]. *k*ache-hash’s keys do not: all the *k*-mers of a minimizer share its buckets. We thus bound the costs per run, whatever the skew. Appendix B.1 also shows that lookups are exact, despite their early exits (§2.3).

We count the cache lines that operations bring in from memory, as in the external-memory model [1]. A line counts regardless of whether a demand miss or a prefetch brings it in, as in the roofline analysis (§4.6). Most of them arrive by prefetch, so demand misses are fewer (§4.2). We assume that the cache does not evict the lines that a run touches until the run ends. The lookahead (§2.6) keeps at most a few hundred lines in flight, well within a core’s L1 and L2 caches.

### Theorem 1

**(Lines per run)**

*Let a run of k-mers have the buckets* b_0_, b_1_, *and* b_2_. *Its operations and their prefetches bring in at most (a) the lines of* b_0_, *5 in a set, (b) plus the lines of* b_1_ *and* b_2_, *10 in a set, if* b_0_ *is full or fills during the run, (c) plus O* (1) *lines of the overflow table in expectation*.

*Proof*. All the *k*-mers of a run share their buckets (§2.2). So the run’s operations and their prefetches may touch only these three buckets and the overflow table. A set’s bucket spans 5 lines: its header, and 4 lines of 8 keys each. A map’s bucket spans more, with its values (9 lines for 8-byte values). Operations go past b_0_ only once it is full, and the cache keeps each line. In the linear-probing overflow table, a *k*-mer probes O(1) slots in expectation [22].

### Corollary 1

**(Lines per *k*-mer)**

*Let the* N *k-mers of a sequence form* R *runs, none of which spills. Then in a set, operations on the k-mers bring in at most* 5R *lines. In a long random sequence, that is about* 10/ (w + 1) *lines per k-mer, at most*.

*Proof*. The first claim sums Theorem 1(a) over the R runs. Two consecutive *k*-mers span w + 1 ℓ-mers. In a random sequence, these are distinct with high probability.

The least of their hashes is then equally likely at each of the w + 1 positions. The two *k*-mers have different minimizers only if it is at the first or the last position. So a run starts at a *k*-mer with probability about 2 / (w + 1) at most [43]. With linearity of expectations, the sequence has R = 2N/(w + 1) runs. They bring in 5R lines, so lines per *k*-mer is 10/(w + 1).

This is 0.91 lines for k = 31 and ℓ = 22, *k*ache-hash implementation’s default. Tables that place each *k*-mer by its own hash do worse on distinct *k*-mers. Each *k*-mer’s slot then lies on a random one of the table’s T lines. With C lines of cache, that line is cached with probability at most C/ T. Such a table brings in at least 1 − C /T ≈1 line per *k*-mer for tables far exceeding the cache in size, and more if a probe reads some metadata apart from the slots. On the genome reference (§4.1), *k*ache-hash’s builds and its positive and mixed queries bring in 0.84–1.02 lines per *k*-mer (§4.6). These counts include spills, which Corollary 1 leaves out. At the evaluation’s load factor α = 0.8, 16.0–19.2% of the keys are outside their primary buckets (Appendix Table 3), which fall in the cases left by Corollary 1. The other dynamic tables bring in 1.5–5.1 lines per *k*-mer.

Typical runs bring in fewer lines than Theorem 1 allows. A run’s new *k*-mers fill consecutive slots, so not all lines of a bucket are necessarily touched. And an absent *k*-mer reads a slot only on a false checksum match (§2.1).

### Spills and resizes

Appendix B.2 bounds what spills and resizes add. Even if every run spilled, operations would bring in *O* (1) lines per *k*-mer in expectation. The spill filter’s false-positive rate is at most c/ 254 for a primary bucket shared with c other minimizers, about 1.8% for the average c. On average, resizes move a key at most about twice.

## 4 Evaluation

We evaluate *k*ache-hash against six other tables on five indexes (§4.1), which answer five questions:

§4.2 *How fast does* `kache-hash` *build k-mer tables?* 3.5– 7.0 × faster than the fastest baseline on every index, presized or resizing.

§4.3 *How fast does it answer queries?* 1.5–5.4× faster, with all, almost none, or some of their *k*-mers present.

§4.4 *How does it scale with threads?* From 1 thread to 56, its builds speed up 34–37×, and its queries 29–43×.

§4.5 *How fast is it on one thread?* 1.8–4.9× faster than the fastest baseline.

§4.6 *What limits it?* Memory bandwidth: its builds and positive and mixed queries reach 77–85 % of the band-width for random access, while its negative queries are bound by instruction issue.

We validate *k*ache-hash against an exact reference, the sorted *k*-mers of its input with their counts. In every run below, it also agrees with the other tables on the distinct *k*-mers and on every query’s hits.^5^ The appendix explores the impact of the minimizer size (Appendix C.1), and how keys distribute over the table’s buckets (Appendix C.2).

### 4.1 Experimental Setup

#### Platform

We run every experiment on a server with two Intel Xeon Platinum 8276 processors. These are Cascade Lake processors with 28 cores each, at 2.2 GHz and up to 4.0 GHz with Turbo Boost, supporting AVX-512, with 32 KB of L1d and 1 MB of L2 cache per core. The server has 38.5 MB of L3 cache per socket, and 187 GB of DRAM over two NUMA nodes, under Linux 5.14 (RHEL 9.3) with transparent huge pages always on. With all its cores, it reads DRAM at 128 GB s sequentially and at 104 GB s in random cache lines (§4.6). Every reported run had exclusive use of the server. A run on t threads pins them to one core each, alternating between the two sockets, and interleaves its memory over both NUMA nodes. All code is compiled with GCC 11.5 (`-O3 -march=native`) and allocates through `jemalloc 5.3.0`.

#### Datasets

We index five datasets, each representative of a setting in which *k*-mer tables are used in practice (Table 1). Together they span the main parameters that affect *k*-mer table performance: its size, from 0.17 to 2.5 billion distinct *k*-mers; the redundancy of its input, from 0.9% of the *k*-mers distinct to 81%; and the length of the input’s fragments, which bounds the runs of consecutive *k*-mers that share a minimizer, from 150-bp reads to whole chromosomes. The *reference* is T2T-CHM13v2.0, the complete human genome [35]. The *short reads* are the first 20 M read pairs (2 × 150 bp, 1.9 × coverage) of an Illumina NovaSeq X run of the GIAB sample HG002 [51] (SRA SRR37356339), all first mates before all second mates. The *long reads* are the first 360 K reads (1.5 coverage) of a PacBio HiFi run of HG002 (SRR10382246). The *pangenome* is a public collection of 3,682 *E. coli* assemblies, used in other index evaluations [12,2]. The *metagenome* is every 20th genome of the HumGut collection of human gut genomes [15], also used in various evaluations [20,12,21].

**Table 1.**
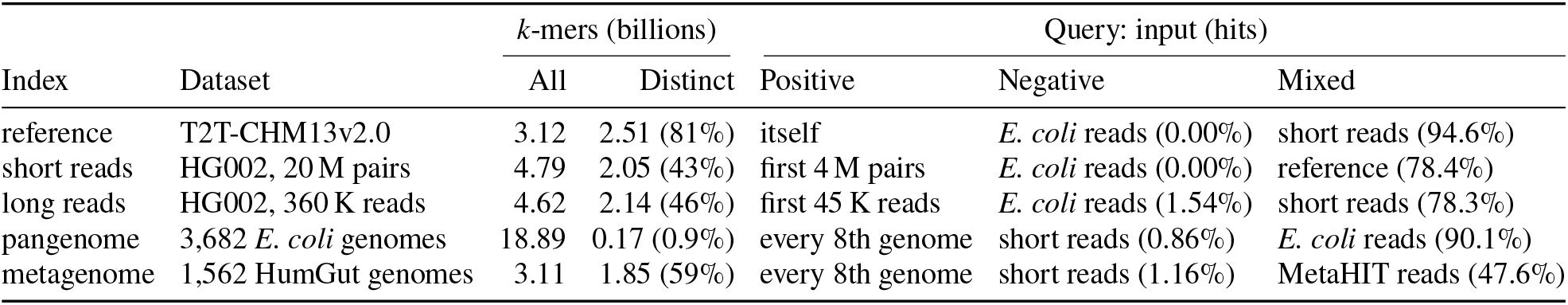
The indexes and their queries. A query’s hits are the share of its *k*-mers present in the index; a positive query’s are all present. The positive queries hold 3.12, 0.96, 0.58, 2.35 and 0.39 billion *k*-mers, in the table’s order. The other query inputs are the *E. coli* reads (SRR5901135; 0.79 billion *k*-mers), the first 4 M pairs of the short reads (0.96), the reference (3.12), and the MetaHIT reads (ERR321482; 0.82).

|  |  | *k*-mers (billions) |  | Query: input (hits) |  |  |
|---|---|---|---|---|---|---|
| Index | Dataset | All | Distinct | Positive | Negative | Mixed |
| reference | T2T-CHM13v2.0 | 3.12 | 2.51 (81%) | itself | *E. coli* reads (0.00%) | short reads (94.6%) |
| short reads | HG002, 20 M pairs | 4.79 | 2.05 (43%) | first 4 M pairs | *E. coli* reads (0.00%) | reference (78.4%) |
| long reads | HG002, 360 K reads | 4.62 | 2.14 (46%) | first 45 K reads | *E. coli* reads (1.54%) | short reads (78.3%) |
| pangenome | 3,682 *E. coli* genomes | 18.89 | 0.17 (0.9%) | every 8th genome | short reads (0.86%) | *E. coli* reads (90.1%) |
| metagenome | 1,562 HumGut genomes | 3.11 | 1.85 (59%) | every 8th genome | short reads (1.16%) | MetaHIT reads (47.6%) |

Each index is evaluated on three categories of queries, each looking up every *k*-mer occurrence of an input. A *positive* query’s *k*-mers are all present: the reference queries itself, the read indexes a prefix of their own reads, and the bacterial indexes every 8th of their genomes. A *negative* query’s *k*-mers are almost all absent: *E. coli* isolate reads (SRR5901135) for the human indexes, and HG002’s short reads for the bacterial ones. A *mixed* query is a realistic representation of looking up *k*-mers in a table build on related but distinct sequence, hitting 48–95% of its *k*-mers: HG002’s short reads against the reference and the long reads, the reference against the short reads, the *E. coli* reads against the pangenome, and gut metagenomic reads from MetaHIT [23] (ERR321482) against the metagenome.

#### Baselines

We compare *k*ache-hash (v0.7.0) against six tables (Table 2). Three are general-purpose concurrent hash tables, given each *k*-mer’s canonical 2-bit encoding as a 64-bit key from a rolling loop: (1) `Boost`’s concurrent flat tables [7] (1.89.0), whose queries use its bulk lookups; (2) `libcuckoo` [26] (0.3.1); and (3) `IcebergHT` [36] (commit 96b9d34), which stores sets only. Three are *k*-mer-specific, and receive the sequences directly: (1)`Jellyfish`’s lock-free counting table [30] (2.3.1); (2)`CBL` [32] (commit 986d57e), a dynamic but single-threaded *k*-mer set; and (3) `SSHash` [39] (6.0.0), a static *k*-mer dictionary with concurrent queries. At one thread, *k*ache-hash and `Boost` run their sequential tables.

**Table 2.** The tables compared: whether each allows insertions (dynamic), runs its operations concurrently, and counts (maps), and whether it is specific to *k*-mers, taking sequences rather than *k*-mers.

| Table | Dynamic | Concurrent | Maps | *k*-mer-specific |
|---|---|---|---|---|
| *k*ache-hash | ✓ | ✓ | ✓ | ✓ |
| Boost | ✓ | ✓ | ✓ |  |
| libcuckoo | ✓ | ✓ | ✓ |  |
| IcebergHT | ✓ | ✓ |  |  |
| Jellyfish | ✓ | ✓ | ✓ | ✓ |
| CBL | ✓ |  |  | ✓ |
| SSHash |  | queries |  | ✓ |

#### Measurements

Each run for the dynamic tables first parses the index’s input into memory on one thread, untimed. Sequences are split at every character other than A, C, G and T (a common strategy in computational genomics tools dealing with *k*-mers that discard ambiguous nucleotides). k is set to 31 throughout. The run then times the table’s construction, which entails inserting every *k*-mer occurrence of the input (a map counts them, while sets maintain no count). It then times each query, which looks up every *k*-mer occurrence of the query’s input in the index. Throughput is in millions of *k*-mer operations per second (M ops/s). Each thread takes a contiguous share of the input. A table is either presized for the input’s distinct *k*-mers, at the table’s own maximum load factor, or sized to half the *k*-mers, so that it resizes once. Runs use all 56 cores unless noted. We report the median of three repetitions, with whiskers to their minimum and maximum. Each repetition also counts one group of hardware events, which has no measurable cost. Memory is the build’s peak resident set, minus the size of the parsed input. Across all runs for the dynamic tables, every run agrees on its distinct *k*-mers and on every query’s hits across the indexes, but for `IcebergHT`’s resized builds, which lose and duplicate keys. The distinct counts also equal those reported by `SSHash` for the corresponding datasets.

### 4.2 Build Performance

*k*ache-hash builds *k*-mer sets at 863–1894 M ops/ s when presized, 3.5–5.8× faster than the fastest baseline on each index (Fig. 2a). The fastest baseline on 3 of the 5 datasets is `Jellyfish`. *k*ache-hash’s builds incur 0.010–0.063 demand L3 misses per *k*-mer, against 0.89–1.99 for the other tables (Appendix Fig. 10), as consecutive *k*-mers tend to share a bucket. *k*ache-hash also prefetches its secondary buckets ahead of their use, and these counts leave out prefetched cache lines, as prefetches do not count as demand misses. The roofline analysis (§4.6) counts every line fetched.

**Fig. 2.**
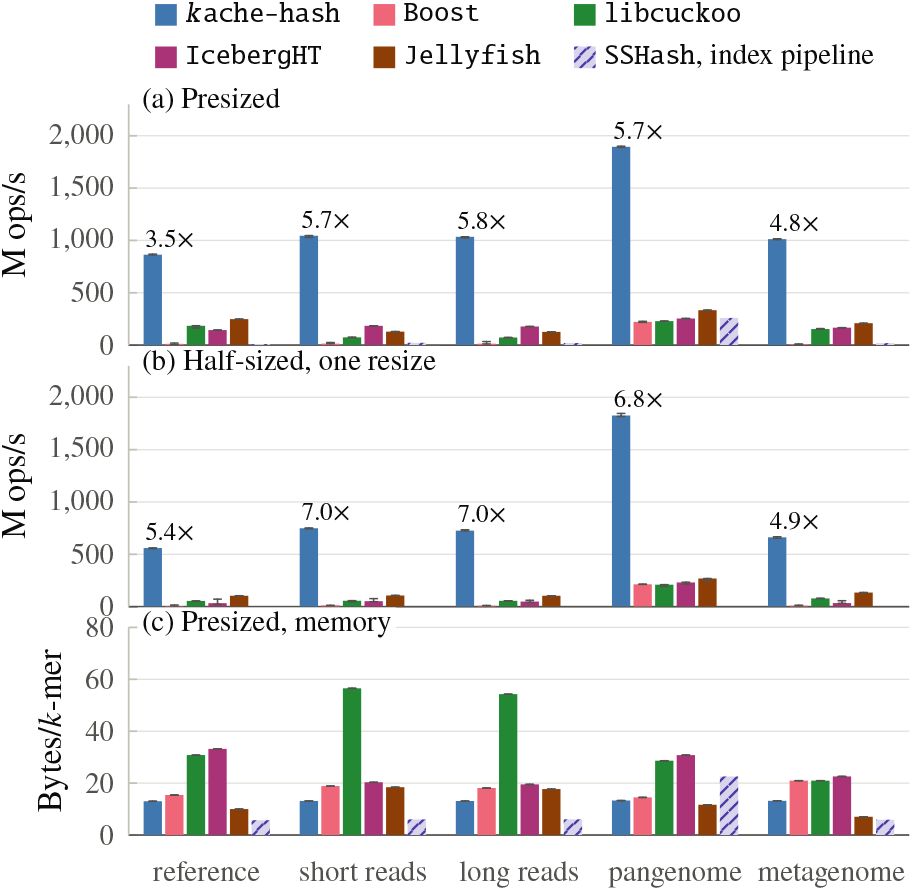
Builds at 56 threads: (a) throughput of tables presized for the index’s distinct *k*-mers, (b) of tables sized for half of them, which resize once, and (c) the presized tables’ memory per distinct *k*-mer. Bars are the median of three runs, with whiskers to the minimum and maximum. Above *k*ache-hash’s bars in (a) and (b) is its lead over the fastest baseline on that index. The hatched bars are `SSHash`’s indexing pipeline: compacted de Bruijn graph, followed by the index. Its time includes reading the input, and its memory is its peak.

Sized for half of the distinct *k*-mers, every table resizes once (Fig. 2b). The resize costs *k*ache-hash 4–35 % of its presized rate, against 9–79 % for `libcuckoo`, `IcebergHT` and Jellyfish. *k* ache-hash still leads on every index, by 4.9–7.0× over `Jellyfish`.

`SSHash`’s indexing pipeline (hatched) runs at 10–259 M ops / s, timed from reading its input, which the dynamic tables’ runs parse untimed on one thread. `SSHash`’s indexing pipeline, from raw input sequences to the final *k*-mer set, cannot receive in-memory sequences as the harness of the dynamic tables does.

Maps, which count their *k*-mers, move alike (Appendix Fig. 11): *k*ache-hash’s maps build 10–15 % slower than its sets, and lead the other maps by 3.4–7.4× presized and 4.2 –6.6× half-sized.

Fig. 2c shows the presized sets’ memory. *k*ache-hash’s sets take 13.0–13.3 bytes per distinct *k*-mer, against `Boost`’s 14–21, `Jellyfish`’s 7.0–18.4, `IcebergHT`’s 20– 33 and `libcuckoo`’s 21–57 (Appendix Table 4). Its tables end 80% full on every index, at 10 bytes per slot, beside a small overflow table. The other tables round their capacities up to powers of two, so they end 46–89% full. Among the dynamic tables, only `Jellyfish` takes less than *k*ache-hash on three datasets, on the reference, the pangenome, and the metagenome. It truncates its keys by quotienting. `SSHash`’s index takes just 1.2–1.4 bytes per *k*-mer, but its pipeline (hatched) peaks at 5.7–22.5.

### 4.3 Query Performance

*k*ache-hash answers positive queries on its presized *k*-mer sets at 1394–2886 M ops/ s, 2.2–5.4× faster than the fastest baseline on each index (Fig. 3a). The fastest baseline is `SSHash` on the read indexes and the metagenome, and `Boost` on the reference and the pangenome. On mixed queries, *k*ache-hash runs at 1517–2452 M ops /s, 2.1– 3.8 × faster than `Boost`, the fastest baseline on every index (Fig. 3c). Negative queries are the fastest for both: *k*ache-hash answers them at 2679–3016× M ops /s, 1.5–1.8× faster than `Boost` (Fig. 3b). Across the three kinds of query, *k*ache-hash leads `SSHash`’s static index by 2.2–8.0×, and `libcuckoo`, `IcebergHT` and `Jellyfish` by 3.5–29 × (Appendix Table 5).

**Fig. 3.**
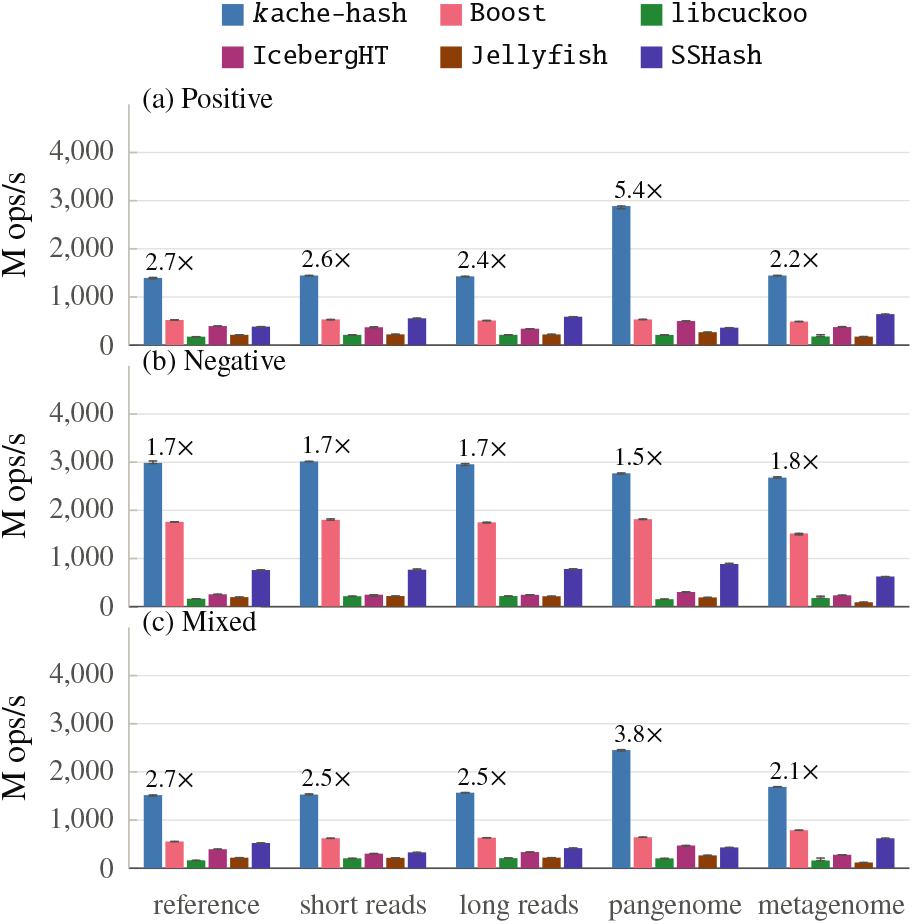
Query throughput at 56 threads on presized tables:(a)positive queries, every *k*-mer present; (b) negative queries, almost none present; and (c) mixed queries (Table 1). Bars are the median of three runs, with whiskers to the minimum and maximum. Above *k*ache-hash’s bar is its lead over the fastest baseline on that index.

*k*ache-hash’s queries incur 0.009–0.100 demand L3 misses per *k*-mer, against 0.16–2.54 for `libcuckoo`’s, `IcebergHT`’s and `SSHash`’s (Appendix Fig. 12). `Boost`’s bulk lookups and Jellyfish’s lookups prefetch the table slots they probe, and so incur only 0.002–0.037, in the order of *k*ache-hash’s.

Maps perform similarly (Appendix Fig. 13): *k*ache-hash maps answer queries 2–11 % slower than its sets, and lead the fastest baseline, `Boost`, by 1.5–4.9×.

### 4.4 Concurrency Scaling

*k*ache-hash’s presized builds on the reference run 36.4× faster on 56 threads than on 1, and its queries 34–41× faster (Fig. 4). It leads every table at every thread count: by 3.5– 4.5× on presized builds, 3.1–6.9× on half-sized builds, and 1.5–2.8× on queries. `libcuckoo`’s, `IcebergHT`’s and `Jellyfish`’s presized builds scale 35–45×, but from 3.8–5.5 M ops/s on one thread, against *k*ache-hash’s 24 M ops/s; `Boost`’s peak at 16 threads. The other tables’ queries scale 20–48×.

**Fig. 4.**
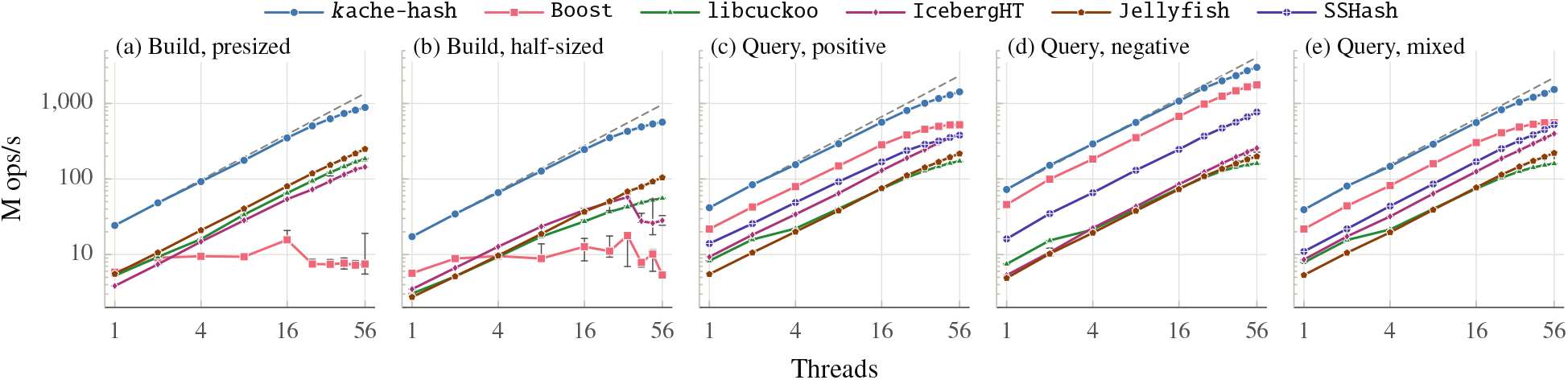
Throughput on the reference from 1 to 56 threads on logarithmic axes: builds of (a) presized tables and (b) tables sized for half of the distinct *k*-mers, which resize once; and (c) positive, (d) negative, and (e) mixed queries on presized tables. Points are the median of three runs, with whiskers to the minimum and maximum. The dashed line is *k*ache-hash’s perfect (linear) speedup sketch.

*k*ache-hash’s speedup is linear in CPU cycles: its build takes 163.4 cycles per *k*-mer on one thread and 158.9 on 56, and incurs 0.061–0.067 demand L3 misses per *k*-mer at every count (Appendix Fig. 14). However, this does not translate to perfect linear scaling of 56 in wall time, due to the CPU clock and idle time. The server’s (§4.1) Turbo Boost runs a lone core at 4.0 GHz, but the build’s 56× cores at 2.79 GHz. And threads that finish their share of the input early wait for the rest, so the cores are busy for 91 % of the build. Together, they account for the measured speedup within 1 %. It scales alike on all the other indexes: 34–37× on presized builds, and 29–43× on queries.

*k*ache-hash’s half-sized builds scale 32.7× (Fig. 4b) on the reference: its resize adds 52 s to the build on one thread, and 2.0 s on 56. The other tables’ resizes do not scale as well. `Jellyfish`’s builds scale 38.2×, but from 2.7 M ops /s on one thread, where its resize takes longer than its whole presized build.

`Boost`’s positive and mixed queries level off from 40 threads, as they approach the memory’s bandwidth for random cache lines (§4.6): from 40 threads to 56, they gain 4–5 %, against *k*ache-hash’s 23–27 %. So *k*ache-hash’s lead over `Boost` grows from 1.8–1.9× on one thread to 2.7–2.8× on 56, while it holds at 1.5–1.7× on negative queries.

### 4.5 Sequential Performance

On one thread, *k*ache-hash’s sequential table builds *k*-mer sets at 28–65 M ops /s, 3.1–4.9× faster than `Boost`’s, the fastest baseline on every index, and 13–14 × faster than `CBL` (Fig. 5a). It answers queries at 40–86 M ops/s, 1.8–3.6× faster than the fastest baseline on each index (Fig. 5b–d), `SSHash` on the metagenome’s positive query and `Boost` on the rest. *k*ache-hash’s queries lead CBL’s by 14–30×, and `SSHash`’s by 2.3–11× (Appendix Table 6).

**Fig. 5.**
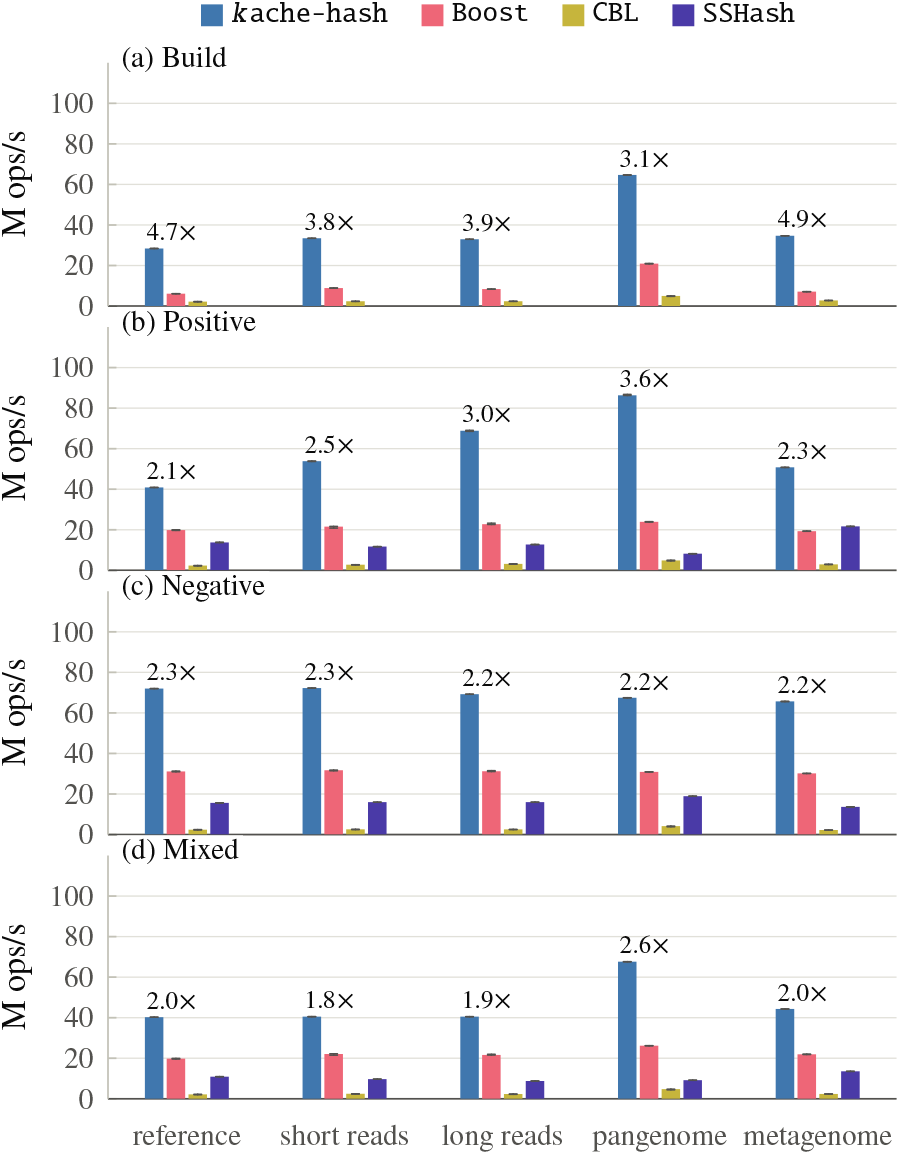
Throughput on one thread: (a) builds, and (b) positive, (c) negative, and (d) mixed queries (Table 1). *k*ache-hash and `Boost` run their sequential tables, presized for the index’s distinct *k*-mers; `SSHash` queries its index built on 56 threads. Bars are the median of three runs, with whiskers to the minimum and maximum. Above *k*ache-hash’s bar is its lead over the fastest baseline on that index.

### 4.6 Roofline Analysis

The roofline model [49] bounds a program’s performance by the limiting hardware resources, specifically, peak compute rate and memory bandwidth. Since *k*-mer tables perform no floating-point arithmetic, we count instructions instead of FLOPs, similar to instruction roofline analysis [9] (Fig. 6). The compute roof is the 56 cores’ peak of 4 instructions per cycle at 3.0 GHz, 672 billion per second: Turbo Boost runs all 56 cores at 2.6–3.1 GHz across the tables’ runs, against a lone core’s 4.0 GHz (§4.1). The memory roofs are the server’s bandwidths, which are 128 GB/s for sequential reads and, for random cache lines, 104 GB/s for reads and 66 GB/s for updates, where lines are also written back. For each table, the analysis takes the numbers of instructions executed and cache lines fetched from memory, prefetching included.

**Fig. 6.**
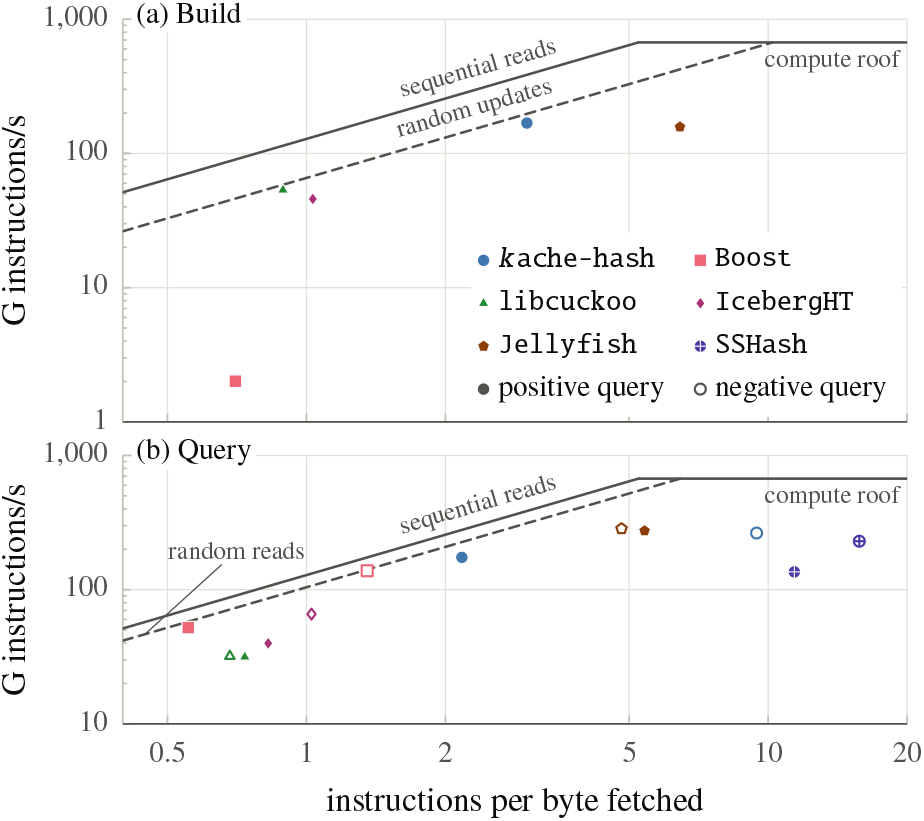
Instruction roofline on the reference at 56 threads, on logarithmic axes: (a) builds, and (b) positive (filled) and negative (hollow) queries, on presized tables. A point’s height is the instructions that the 56 cores retire per second. Its position on the x-axis is the instructions it executes per byte fetched from beyond the L3 cache, prefetched lines included. The solid roofs are the server’s hardware limits: the cores’ peak of 4 instructions per cycle at an all-core clock of 3.0 GHz (flat) and the bandwidth of sequential reads (slanted). The dashed roof is the bandwidth of probing random cache lines, probes being updates for builds and reads for queries. Heights count instructions, not *k*-mers: a table’s *k*-mer rate is its height over its instructions per *k*-mer. Appendix Table 7 lists every point, mixed queries included.

*k*ache-hash’s builds and its positive and mixed queries run at 77–85 % of the random-access roof, so memory bandwidth bounds them. It fetches the fewest cache lines per *k*-mer of the dynamic tables: 0.84–1.02, against 1.5–5.1 for the others. Its negative queries fetch only 0.15 lines per *k*-mer, and are bound by instruction issue instead, at 39 % of the compute roof. `Boost`’s queries also run at the roof (90–98 %), but fetch 2.7–2.8 lines per *k*-mer on positive and mixed queries. Its builds reach only 4 % of the roof, stalled on memory for 99 % of their cycles but on DRAM for only 5 %, likely a sign of contention. `libcuckoo`’s and `IcebergHT`’s queries reach 39–62 % of the roof, stalled for 68–82 % of their cycles waiting on cache misses. `Jellyfish`’s queries reach 49–57 %: it fetches 3.7–4.7 lines and runs 1290–1440 instructions per *k*-mer, 10–16× *k*ache-hash’s, which is why its points sit high. `SSHash`’s queries are bound by instruction issue, reaching 20–34 % of the compute roof.

## 5 Applications Demonstration

Section 4 measured *k*ache-hash’s performance in isolation. Here we measure its effect on two existing tools built for different purposes, `PanGenie` [10] and `K2Rmini` [34].

*k*ache-hash is generic and easy to adopt, and the results suggest what other tools with dynamic *k*-mer tables can expect.

### Experimental setup

Each demonstration replaces the *k*-mer table of an application’s latest release with *k*ache-hash, parallelizes the steps around the table that would otherwise bound it, and leaves the rest as released. We time the application’s own command end to end, report the median of three runs, and check every run’s output against the release’s. We evaluate the application workloads on a second server, since the primary server (§4.1) has 187 GB of DRAM, and `PanGenie`’s reads alone take 164 GB. The server has two 16-core Intel Xeon Gold 5218 processors (Cascade Lake, as in §4.1) and 754 GB of DRAM, under Linux 5.14 and GCC 11.5.

#### 5.1 Targeted counting for pangenome genotyping

`PanGenie` [10] genotypes a newly sequenced individual against a pangenome, a panel of haplotype-resolved genomes. At each site where they differ, it infers which alleles the individual carries from how often each allele’s *k*-mers occur in its short reads, without aligning them. It counts every *k*-mer of the panel’s graph (k = 31) in the reads, estimates their coverage from the counts’ histogram, then looks up each site’s allele-specific *k*-mers and infers genotypes with a hidden Markov model. It has been used to genotype all 3,202 samples of the 1000 Genomes Project (1kGP) this way [11]. It uses `Jellyfish` [30] for its *k*-mer counting.

### The workload

Counting streams sequences against a *k*-mer table far larger than the caches, and performs the following operations. (1) *Priming* builds the table from the graph’s sequences, 3.58 billion distinct *k*-mers, so that only these are counted in the reads. (2) *Counting the reads* increments the counts of the *k*-mers from the reads that are present in the table, 88.0 billion, and skips the rest. (3) *The histogram* scans the table to extract the final counts (on one thread as released). The later steps in `PanGenie` look up single *k*-mers. We swap *k*ache-hash in for `Jellyfish` in `PanGenie`’s latest release, v4.2.2 [19].

### Setup

The panel is HPRC’s v2.0 pangenome [16] as distributed by `PanGenie` [11]: 462 haplotypes on CHM13 [35] with 23.4 million variant sites. The sample is HG00096, outside the panel: 1kGP’s 37× NovaSeq run [8], 768 million reads in one uncompressed FASTA file. `PanGenie`’s 1kGP pipeline also takes its reads as FASTA, gzipped. `PanGenie` runs its 1kGP command with 24 threads; Appendix D.2 varies the counting threads from 8 to 32. A control keeps `Jellyfish`’s table in the backend, behind *k*ache-hash’s parallel reader and parallel histogram scan, so that it differs from *k*ache-hash only in the table. All runs, with `Jellyfish`, the control, and *k*ache-hash, output the same VCF and histogram.

### Results

With *k*ache-hash, `PanGenie` genotypes the sample 1.96 faster, in 1976 s instead of 3876 s (Fig. 7). Counting, which is 54% of the run as released, runs

**Fig. 7.**
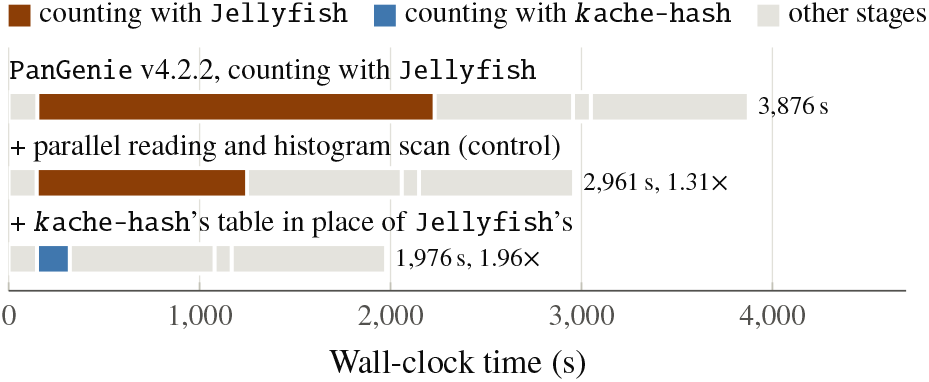
PanGenie genotyping HG00096 by stage: as released, with the control, and with *k*ache-hash, along with their speedups. All are medians of three runs.

12.1× faster, falling to only 9% of the total runtime. Of the counting time saved, 50% is *k*ache-hash’s table, which primes 3.84× and counts the reads 7.38× faster than `Jellyfish`’s behind the same reader, 35% the histogram scan, which `PanGenie` runs on one thread, and 15% the parallel reader. Against the control, *k*ache-hash counts 6.42× faster, and genotypes 1.50× faster. The rest of the run is mostly `PanGenie`’s own work, including its single-threaded VCF writing and the haplotype sampling.

*k*ache-hash also reduces the CPU time per sample from 12.3 to 4.2 CPU-hours: about 26,000 CPU-hours over 1kGP’s 3,202 samples. But it takes more memory: a 90.2 GB peak against 51.3 GB, as its table keeps keys whole (63.5 GB) where `Jellyfish` truncates keys by quotienting (24.2 GB). However, both still stay under the 106 GB that building `PanGenie`’s index takes, so that the peak RSS of the entire pipeline is unchanged. `PanGenie`’s 1kGP pipeline reads its gzip files through a single-threaded `gunzip`, and decompression would bound *k*ache-hash’s counting; a parallel decompressor recovers most of the gain (Appendix D.3).

#### 5.2 Read filtering by *k*-mer content

`K2Rmini` [34] keeps every read in which at least a fraction t of the *k*-mer instances occur in a set of patterns, such as a host genome whose reads should be depleted or a species’ pangenome whose reads should be recruited. It rules most reads out by their *k*-mers’ minimizers, and then counts the *k*-mers (k = 31) of the rest in a hash set of the patterns’ *k*-mers. The authors identify that set as `K2Rmini`’s main limitation, and propose a concurrent table that groups *k*-mers by their minimizers in its place [34].

### The workload

At genome scale, the set is far larger than the caches. (1) *Indexing* inserts every *k*-mer instance of the patterns, 6.2 billion for the human genome with its reverse complement, on one thread. (2) *Filtering* looks up every *k*-mer of each read that the minimizers do not rule out. `K2Rmini` stores 32-bit hashes of the *k*-mers, so a hash collision counts an absent *k*-mer as present, noted in its documentation [33]. We swap `k ache-hash` in for this set in `K2Rmini`’s latest release, 1.0.0 [18]. We store the exact *k*-mers in canonical form, with a one-byte mask of the strands they occur on, as `K2Rmini` matches forward strands only. Other than the *k*-mer table, we only update `K2Rmini` by reading the patterns in parallel and keeping a sharded minimizer set instead of a monolithic one, so that neither becomes the next serial step. The release’s minimizer set hashes with `FxHash`, which skews heavily in the human genome. Using `Murmur3`’s finalizer instead, the release keeps the same output and runs 4.7–4.9× faster there (Appendix D.4). We compare against this *fixed control*, with the release alongside.

### Setup

Two tasks filter HiFi reads at t = 0.8 against the complete human genome, T2T-CHM13v2.0 [35]: the `K2Rmini` paper’s own HG002 reads (SRR8858457 [48]; 92 K reads), and as a host depletion, an eighth of a saliva sample’s run (SRR32782080 [13]; 1.0 M reads), shuffled out of its genome order. The third uses *E. coli* reads from an infant gut metagenome (ERR3405986 [44]; 16.3 M Illumina reads), as in [28], against the evaluation’s 3,682 *E. coli* genomes (§4.1), at `K2Rmini`’s default t = 0.5. `K2Rmini`’s own benchmark filters the same HG002 reads against up to 2^27^ *k*-mers of the reads themselves (positive patterns) or of a random sequence (random ones). We extend it to 2^28^ positive and 2^30^ random patterns. `K2Rmini` runs its own command with 32 threads.

## Results

With *k*ache-hash, `K2Rmini` runs the three tasks 15.2×, 10.7× and 19.4× faster than the fixed control, in 29.9, 44.3 and 95.8 s instead of 453, 475 and 1854 s, and 74×, 51× and 19× faster than the release (Fig. 8a). Indexing takes 94–99% of the control’s runs, and *k*ache-hash builds the index 18.5× and 36.0× faster. Of the indexing time saved over the control, 77% and 72% is *k*ache-hash’s *k*-mer set, and the rest the parallel reading and the sharded minimizer set. Filtering samples against the human genome takes 7.5 min of indexing with the control, and 24 s with *k*ache-hash. On `K2Rmini`’s own benchmark (Fig. 8b), the gain grows with the patterns: trivial up to 2^22^, then 4.14× at 2^26^ and 9.05× at 2^28^ of the reads’ own *k*-mers, and 12.0× at 2^30^ random ones.

**Fig. 8.**
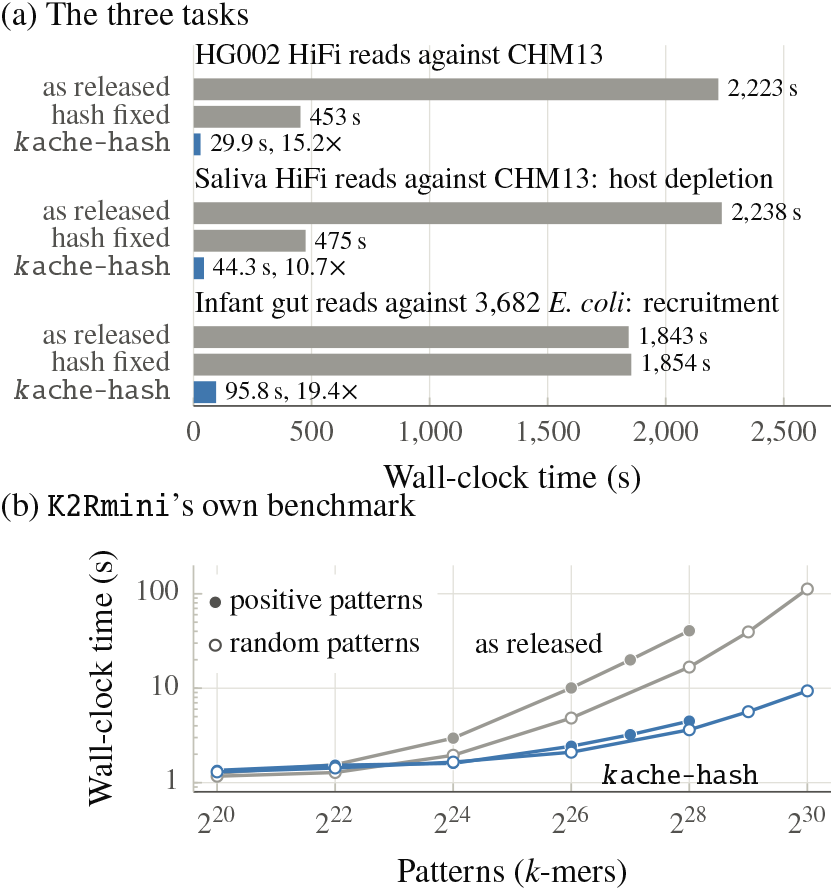
`K2Rmini` as released, with its minimizer set’s hash fixed, and with *k*ache-hash: (a) on the three tasks, with *k*ache-hash’s speedups over the fixed control, and (b) on `K2Rmini`’s own benchmark on logarithmic axes, with positive (filled) and random (hollow) patterns, where the control runs as the release does. Gray marks `K2Rmini`’s own *k*-mer set, and blue *k*ache-hash’s; all are medians of three runs.

*k*ache-hash also makes the filter exact. Against the human genome, the release’s 32-bit hashes of 5.0 billion *k*-mers take 69% of the 2^32^ values, so most absent *k*-mers count as present: the release keeps 97% of the HG002 reads below the threshold (4,133 of 4,267), and 31% of the saliva’s (57,702 of 188,469), which a host depletion would then discard. However, exact keys cost memory: the *k* ache-hash integration has a peak of 82.8 GB against the control’s 56.7 GB on the human genome, and of 42.7 GB against 41.7 GB on the *E. coli* genomes.

## 6 Conclusion

Instead of operating on single *k*-mers, *k*ache-hash makes a run of consecutive *k*-mers with one minimizer the unit of work in a dynamic, concurrent hash table. Each run fetches its buckets once, and an insertion locks them once. A spill filter lets lookups exit early under skew, yet stay exact, and the table grows in place. We show that, under a cache-residency assumption, a run that stays in its primary bucket brings in at most that bucket’s cache lines, however long it is.

In practice, the table fetches at most about one cache line per *k*-mer, and memory bandwidth bounds its builds and most queries. *k*ache-hash builds and queries *k*-mer sets faster than every table we compared, and speeds up genotyping and read filtering end to end. Over `PanGenie`’s 1kGP callset, this would save about 26,000 CPU-hours.

It keeps *k*-mers whole, so it can use more memory than tables that compress or hash their keys. Even so, it demonstrates that sequence locality pays off in fast dynamic tables, not only in compact or static indexes. Storing keys more compactly, without sacrificing throughput, is a natural next step. Minimizer schemes with lower density than random ones would lengthen runs, and may cut the table’s memory traffic further. On the theory side, bounding how often runs spill under realistic minimizer skew remains future work. More broadly, applications that need a dynamic, concurrent *k*-mer table could adopt *k*ache-hash, or its way of sharing cache lines and locks across a run.

## A Design Details

This appendix adds details to the design and implementation of Section 2, from *k*ache-hash’s v0.7.0 code.

### Operations

The operations are (1) `insert`, for a set; (2) `upsert`, for a map, which also applies a function to the values of the present *k*-mers; (3) `update`, which only applies the function; (4) `find` and `count`; (5) and `for_each`, a parallel scan. `upsert` and `find` may also give each *k*-mer’s orientation. Insertions may overlap one another, each thread with its own token. Lookups may overlap one another, and insertions as long as neither table grows meanwhile. A map’s lookup may then read a key’s value as 0, while an insertion is placing the key in the overflow table. Updates may overlap only one another, and a scan no modification.

### Resizing

Each thread adds its new keys to the shared count after at most max(1, t/1024) of them, for the resize threshold t = αmB. With at most 128 threads, the shared count thus lags by fewer than max(128, t/8) keys. At a resize, the new overflow table has room for all the old one’s keys. Most of them overflow again, as a frequent minimizer’s *k*-mers still share their three buckets. A resize’s second phase prefetches in the same two stages as insertions, through a pipeline of 32 keys.

## B Analysis Details

This appendix gives the proofs and details that Section 3 leaves out.

### B.1 Exact Lookups

A lookup may stop at its primary bucket, or before the overflow table (Section 2.3). An invariant of the placement makes these exits safe.

#### Lemma B.1

**(Placement)**. *Let* x *be a key with minimizer* µ. *From* x*’s placement, by an insertion or a resize, until the next resize, one of three cases holds. (i)* x *is in* b_0_. *(ii)* b_0_ *is full and its spill filter marks* µ, *and* x *is in* b_1_ *or* b_2_. *(iii) As in (ii), but* b_1_ *and* b_2_ *are full too, and* x *is in the overflow table*.

*Proof*. An insertion places x past a bucket only once it finds the bucket full, under the bucket’s lock. It marks µ in b_0_’s filter before it places x past b_0_. *k*ache-hash deletes no keys, so until a resize, buckets only gain keys and filters only gain marks. So the case that holds when x is placed stays true until the next resize. A resize clears the filters, and its first phase moves keys only into their new primary buckets. Its second phase inserts the other keys as an insertion would.

#### Theorem B.1

**(Exact lookups)**. *A lookup of a k-mer* x *finds it if and only if* x *is in the table. The lookup probes only* b_0_ *if* b_0_ *has room, or does not mark* µ. *It probes the overflow table only if* b_0_, b_1_ *and* b_2_ *are all full*.

*Proof*. A lookup compares whole keys, so it never finds an absent *k*-mer. Let x be present. If b_0_ has room, or does not mark µ, then x is in b_0_, by Lemma B.1. If b_1_ or b_2_ has room, then x is in one of the three buckets. So no exit misses *x*.

### B.2 Spills and Resizes

How often runs spill depends on how skewed the minimizers’ frequencies are. We provide no bound that holds for every input characteristic. On the five indexes of Section 4, at load α = 0.8, 80.8–84.0% of the keys end up in their primary buckets, 14.3–16.8% in secondary ones, and 1.7–3.8% in the overflow table (Table 3). So Corollary 1’s assumption of no spills holds for most keys.

Theorem 1 bounds the cost of spills. A spilled run adds at most the 10 lines of its secondary buckets, however long the run is. A spilled *k*-mer adds *O* (1) lines of the overflow table in expectation [22], prefetches included. Even if *every* run spilled, operations would thus bring in *O* (1) lines per *k*-mer in expectation, like a linear-probing table, no worse than a general hash table.

Besides crowding the main table, skew also costs lookups of absent *k*-mers, as many primary buckets spill. The spill filter (Section 2.1) keeps this cost low.

#### Lemma B.2

**(Spill filter)**. *Let a k-mer* x *be absent, and let its minimizer* µ *have no k-mers past* b_0_. *Let* b_0_ *be full, and let it be the primary bucket of* c *other minimizers. Then a lookup of* x *probes* b_1_ *and* b_2_ *with probability at most*

*Proof*. A minimizer marks a flag in b_0_’s filter only if it spilled from b_0_, its primary bucket. A minimizer’s primary bucket comes from the low 32 bits of its hash, and its spill flag from bits 32–39, with 0 mapped to 1 since flag 0 is the lock. Under random hashing, flag 1 thus has probability 2 /256, and each other flag 1 /256. In the same bucket, a minimizer λ ≠ µ will share µ’s flag with probability 2 /256 if the flag is 1, and with 1 /256 otherwise. Thus averaged over µ’s hash, λ’s flag is that of µ’s with probability (2/256)^2^ + 254 ×(1/256)^2^ = 258/256^2^ < 1/254 (and at most 2/256, when colliding on the flag 1). A union bound over the c minimizers completes the proof.

At load α, the table holds αB keys per bucket. In a random sequence, each minimizer has one run of about (w + 1) / 2 *k*-mers (Corollary 1). So there are about 2αB / (w + 1) minimizers per bucket. For k = 31, ℓ = 22, and α = 0.8, this gives c ≈ 4.65 on average, and a bound of 1.8%. Full buckets tend to be the primary buckets of more minimizers, so their bounds are somewhat higher.

#### Resizing

A resize’s first phase reads the old buckets and writes the new ones in order, so it streams through memory. Its second phase re-inserts only the keys outside their primary buckets. These constitute 16.0–19.2% of the keys in our evaluations. Doubling also bounds the total work. The resize to m buckets moves the keys that the table then holds, about αBm/2. The resizes before it move fewer keys in all, about αB(m /4 + m /8 + · · ·) < αBm /2. So, on average, a key moves at most about twice.

## C Evaluation Details

This appendix supplements the evaluation of Section 4.

### C.1 Minimizer Size

*k*ache-hash places a *k*-mer by its minimizer, an ℓ-mer (Section 2.2), with ℓ = 22 by default for k = 31. Larger sizes speed up most builds up to ℓ = 24–26, but slow the negative queries past ℓ = 20 (Fig. 9).

**Fig. 9.**
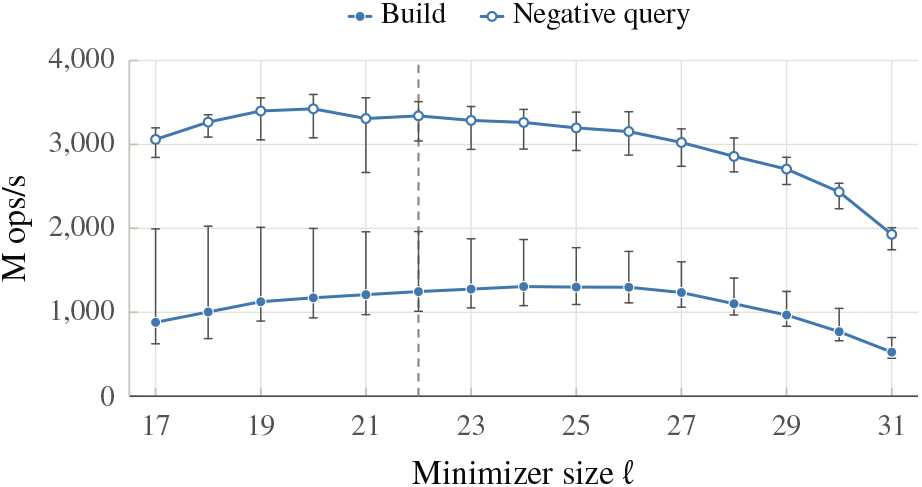
The minimizer sweep: *k*ache-hash’s presized sets, at 56 threads. Throughput of the builds and the negative queries: the median over the indexes, with whiskers to the extremes. The dashed line marks ℓ = 22.

### C.2 Key Distribution

Table 3 shows where *k*ache-hash’s builds leave their keys. The shares hardly depend on the threads: from 1 to 56, they move by at most 0.23 points.

**Table 3.** Shares of *k* ache-hash’s keys in their primary buckets, in their secondary buckets, and in the overflow table, at the ends of the builds at 56 threads (medians of three runs). The tables end 80% full. Maps place their keys as sets do, within 0.01 points.

| Share (%) | reference | short reads | long reads | pan-genome | meta-genome |
|---|---|---|---|---|---|
| Presized sets |  |  |  |  |  |
| primary | 82.0 | 82.0 | 81.7 | 80.8 | 84.0 |
| secondary | 14.3 | 14.3 | 14.5 | 16.8 | 14.3 |
| overflow | 3.7 | 3.7 | 3.8 | 2.4 | 1.7 |
| Half-sized sets |  |  |  |  |  |
| primary | 82.1 | 82.1 | 81.8 | 80.8 | 84.0 |
| secondary | 14.1 | 14.1 | 14.3 | 16.7 | 14.3 |
| overflow | 3.8 | 3.8 | 3.8 | 2.5 | 1.7 |

### C.3 Build Performance

Fig. 10 gives the builds’ demand L3 misses, Fig. 11 the maps’ build throughput, and Table 4 the tables’ memory. Half-sized, *k*ache-hash’s builds still incur 8.0–78× fewer misses per *k*-mer than the next-fewest table. A resize raises *k*ache-hash’s peak memory by 16–26% for sets and 18– 26% for maps, against 43–54% for `Boost`, `libcuckoo` and `Jellyfish` where their presized builds do not resize.

**Fig. 10.**
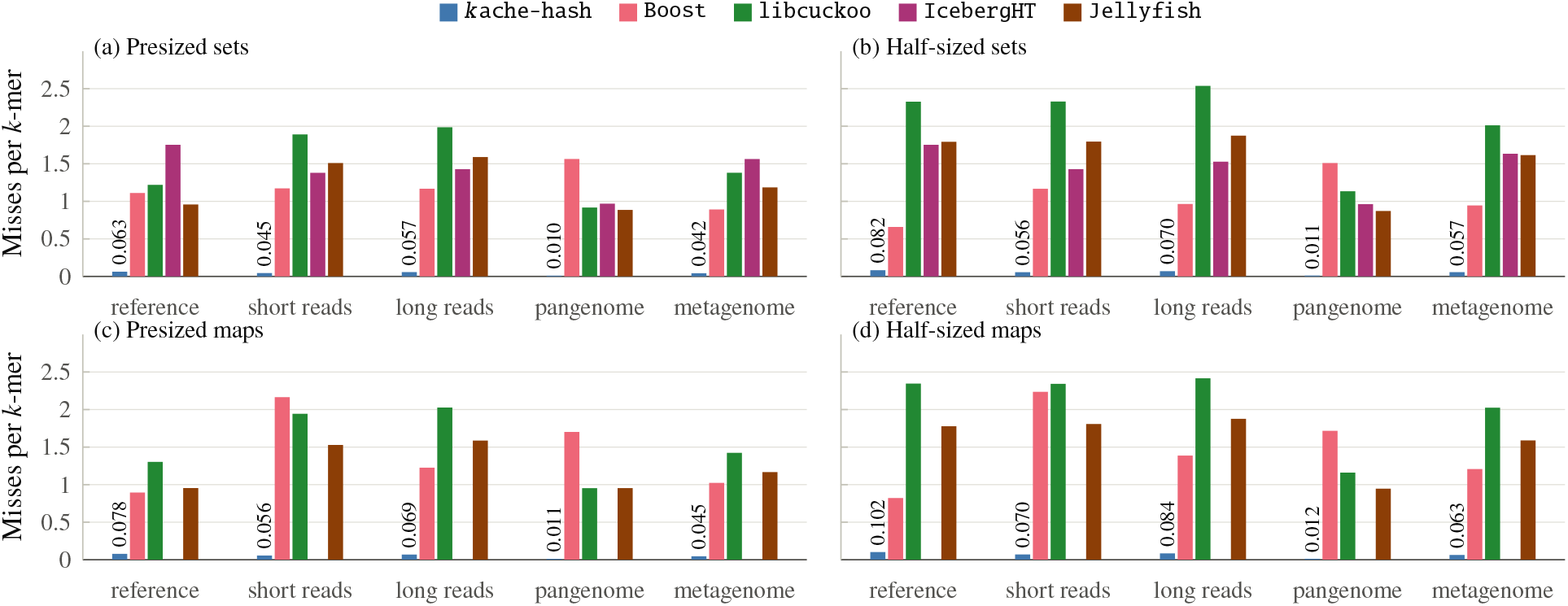
Demand L3 misses per *k*-mer inserted, at 56 threads: (a) presized and (b) half-sized sets, as in Fig. 2, and (c, d) the maps, likewise. Prefetched lines are not counted, which §4.6 counts. Above *k*ache-hash’s bars are their values, since the bars are very short to be perceptible. `IcebergHT` supports sets only, and `SSHash`’s pipeline was not counted.

**Fig. 11.**
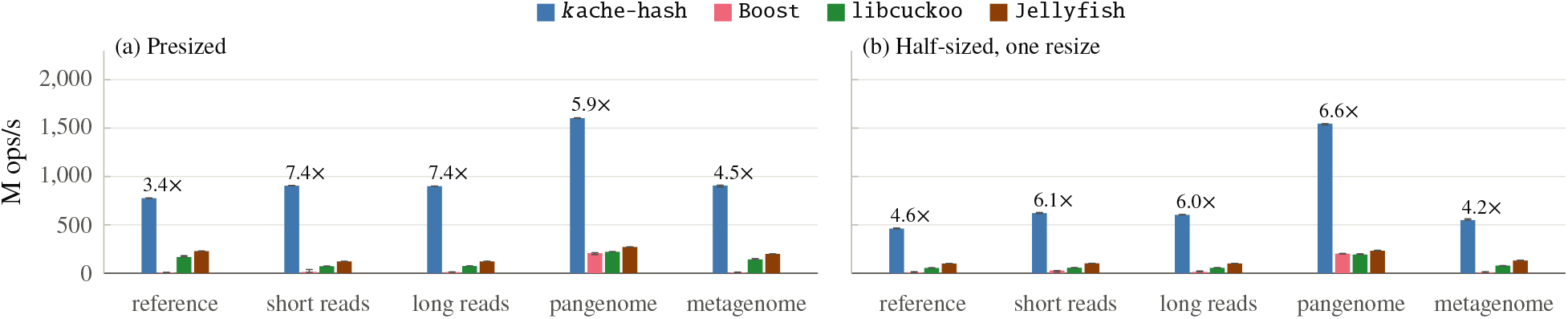
Build throughput of the maps, which count their *k*-mers, at 56 threads: (a) presized, and (b) sized for half the distinct *k*-mers, resizing once. Bars, whiskers and leads are as in Fig. 2. `IcebergHT` and `SSHash` store sets only.

**Table 4.**
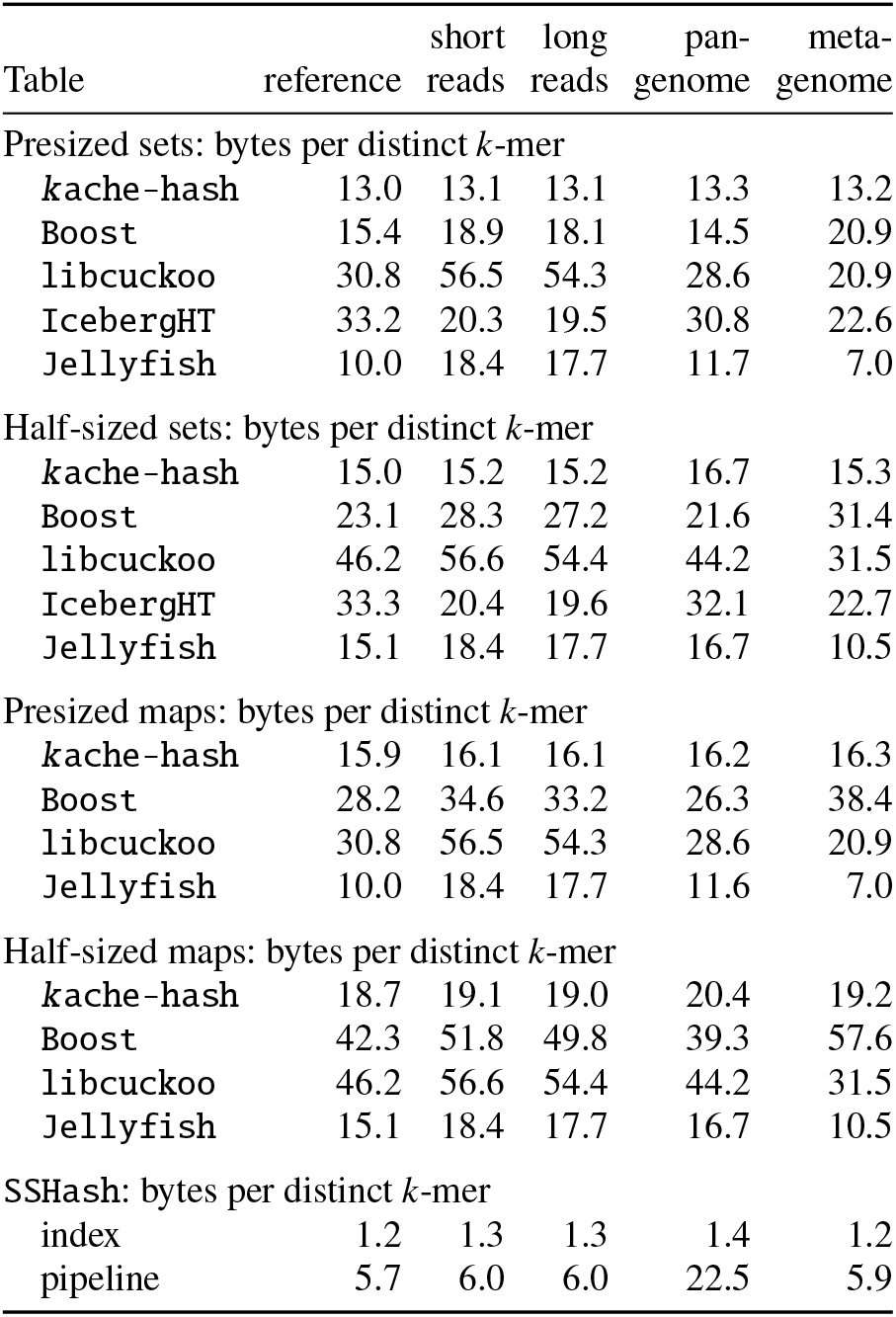
Memory at 56 threads, per distinct *k*-mer of the index: each table’s build peak, minus the parsed input (medians of three runs), and SSHash’s index and its pipeline’s peak. *k*ache-hash’s maps here hold a 1-byte count per key, which wraps past 255, and are from one run. `Boost`’s and `libcuckoo`’s key–count pairs pad to 16 bytes even with 1-byte counts. `Jellyfish` packs a 7-bit count into each entry, so its maps take as much memory as its sets. `IcebergHT` stores sets only.

### C.4 Query Performance

Table 5 gives *k*ache-hash’s query lead over each table, Fig. 12 the queries’ demand L3 misses, and Fig. 13 the maps’ query throughput.

**Fig. 12.**
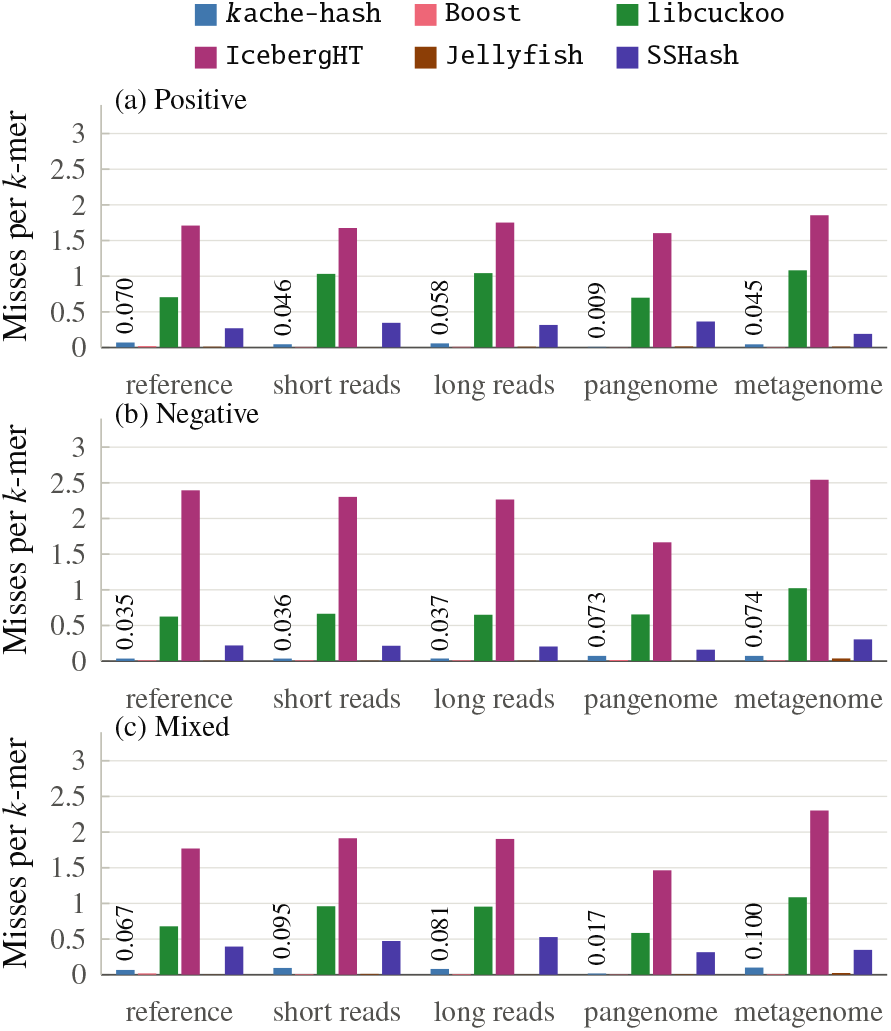
Demand L3 misses per *k*-mer queried, at 56 threads on presized sets: (a) positive, (b) negative and (c) mixed queries, as in Fig. 3. Prefetched lines are not counted. Above *k*ache-hash’s bars are their values. `Boost`’s and `Jellyfish`’s, 0.002–0.037, are also too short to see.

**Fig. 13.**
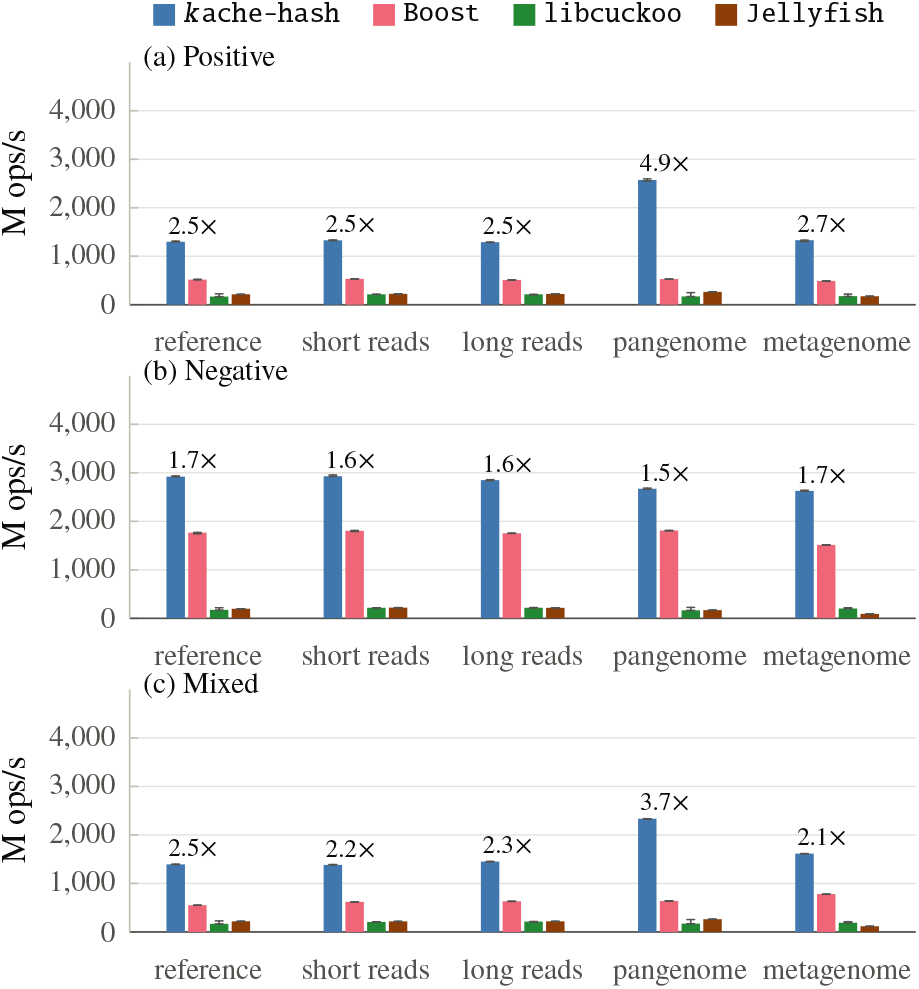
Query throughput of the presized maps at 56 threads: positive, (b) negative and (c) mixed queries. Bars, whiskers and leads are as in Fig. 3, on its y-axis. `IcebergHT` and `SSHash` store sets only.

**Table 5.** *k*ache-hash’s query lead over each other table at 56 threads, on presized tables: the geometric mean over the five indexes of *k*ache-hash’s rate over the table’s, with the minimum and maximum. `IcebergHT` and `SSHash` store sets only.

| *k*ache-hash over | Positive | Negative | Mixed |
|---|---|---|---|
| Presized sets |  |  |  |
| Boost | 3.2 (2.7–5.4) | 1.7 (1.5–1.8) | 2.7 (2.1–3.8) |
| libcuckoo | 8.3 (6.7–14) | 16 (13–18) | 9.2 (7.5–12) |
| IcebergHT | 4.2 (3.5–5.8) | 11 (9.1–12) | 4.9 (3.9–6.1) |
| Jellyfish | 7.5 (6.4–11) | 16 (14–29) | 8.6 (6.9–14) |
| SSHash | 3.3 (2.2–8.0) | 3.8 (3.1–4.3) | 3.8 (2.7–5.7) |
| Presized maps |  |  |  |
| Boost | 2.9 (2.5–4.9) | 1.6 (1.5–1.7) | 2.5 (2.1–3.7) |
| libcuckoo | 8.0 (6.0–15) | 14 (13–17) | 8.5 (6.7–14) |
| Jellyfish | 6.9 (5.8–9.7) | 16 (13–29) | 8.0 (6.3–14) |

### C.5 Concurrency Scaling

Fig. 14 gives the demand L3 misses per *k*-mer at each thread count.

**Fig. 14.**
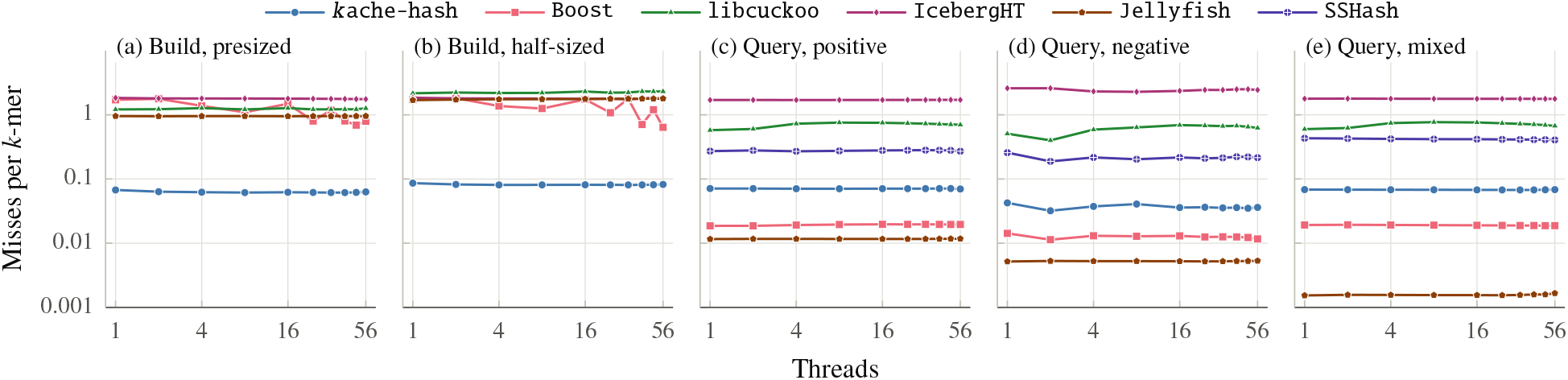
Demand L3 misses per *k*-mer on the reference from 1 to 56 threads, on logarithmic axes, in the runs of Fig. 4: builds of (a) presized and (b) half-sized tables, and (c) positive, (d) negative, and (e) mixed queries. Prefetched lines are not counted.

### C.6 Sequential Performance

Table 6 gives *k*ache-hash’s one-thread lead over each table.

**Table 6.** *k*ache-hash’s lead on one thread over each other table: the geometric mean over the five indexes of *k*ache-hash’s rate over the table’s, with the minimum and maximum. *k* ache-hash and `Boost` run their sequential tables; `SSHash`’s index is built on 56 threads, so it has no build here.

| *k*ache-hash over | Boost | CBL | SSHash |
|---|---|---|---|
| Build | 4.0 (3.1–4.9) | 13 (13–14) | – |
| Positive | 2.7 (2.1–3.6) | 19 (17–22) | 4.5 (2.3–11) |
| Negative | 2.2 (2.2–2.3) | 26 (16–30) | 4.3 (3.6–4.8) |
| Mixed | 2.1 (1.8–2.6) | 17 (14–19) | 4.4 (3.3–7.4) |

### C.7 Roofline Analysis

Table 7 lists every point of Fig. 6, mixed queries included.

**Table 7.**
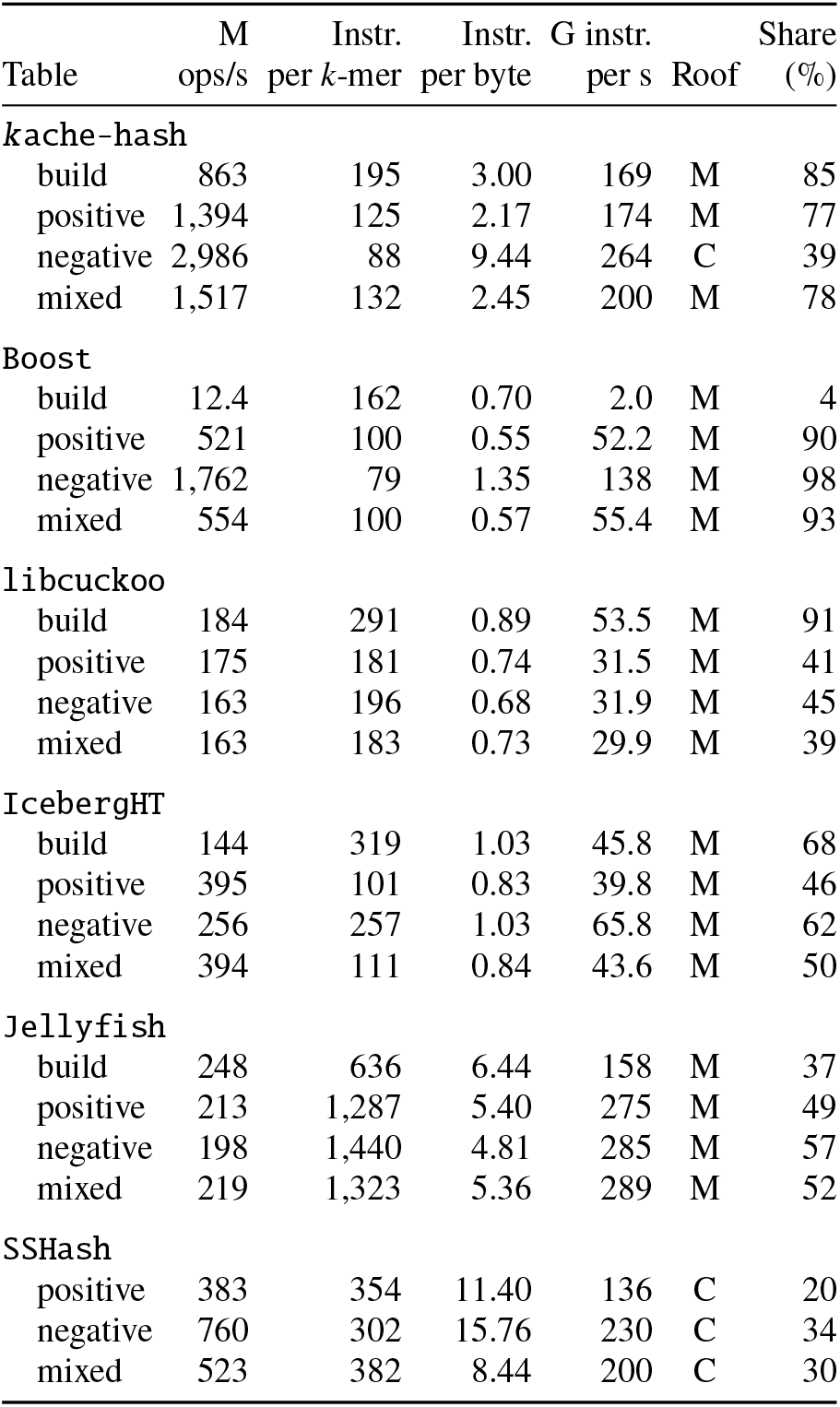
The points of Fig. 6: every table’s build and queries on the reference’s presized sets at 56 threads. A point’s intensity is its instructions per byte fetched from beyond the L3, and its height the instructions retired per second. Its roof is the lower of the compute roof, C, and the memory roof at its intensity, M. The share is its height over that roof.

|  | M | Instr. | Instr. | G instr. |  | Share |
| Table | ops/s | per *k*-mer | per byte | per s | Roof | (%) |
|---|---|---|---|---|---|---|
| *k*ache-hash |  |  |  |  |  |  |
| build | 863 | 195 | 3.00 | 169 | M | 85 |
| positive | 1,394 | 125 | 2.17 | 174 | M | 77 |
| negative | 2,986 | 88 | 9.44 | 264 | C | 39 |
| mixed | 1,517 | 132 | 2.45 | 200 | M | 78 |
| Boost |  |  |  |  |  |  |
| build | 12.4 | 162 | 0.70 | 2.0 | M | 4 |
| positive | 521 | 100 | 0.55 | 52.2 | M | 90 |
| negative | 1,762 | 79 | 1.35 | 138 | M | 98 |
| mixed | 554 | 100 | 0.57 | 55.4 | M | 93 |
| libcuckoo |  |  |  |  |  |  |
| build | 184 | 291 | 0.89 | 53.5 | M | 91 |
| positive | 175 | 181 | 0.74 | 31.5 | M | 41 |
| negative | 163 | 196 | 0.68 | 31.9 | M | 45 |
| mixed | 163 | 183 | 0.73 | 29.9 | M | 39 |
| IcebergHT |  |  |  |  |  |  |
| build | 144 | 319 | 1.03 | 45.8 | M | 68 |
| positive | 395 | 101 | 0.83 | 39.8 | M | 46 |
| negative | 256 | 257 | 1.03 | 65.8 | M | 62 |
| mixed | 394 | 111 | 0.84 | 43.6 | M | 50 |
| Jellyfish |  |  |  |  |  |  |
| build | 248 | 636 | 6.44 | 158 | M | 37 |
| positive | 213 | 1,287 | 5.40 | 275 | M | 49 |
| negative | 198 | 1,440 | 4.81 | 285 | M | 57 |
| mixed | 219 | 1,323 | 5.36 | 289 | M | 52 |
| SSHash |  |  |  |  |  |  |
| positive | 383 | 354 | 11.40 | 136 | C | 20 |
| negative | 760 | 302 | 15.76 | 230 | C | 34 |
| mixed | 523 | 382 | 8.44 | 200 | C | 30 |

## D Demonstration Details

This appendix supplements the demonstrations of Section 5.

### D.1 PanGenie: Integration and Runs

#### Integration

The fork [19] adds a counter to `PanGenie` v4.2.2, chosen by a new option, -`K`, and leaves the rest of `PanGenie` unchanged. The counter maps `PanGenie`’s counting onto *k*ache-hash’s operations: (1) priming onto a batched upsert; (2) counting the reads onto an atomic update of the present *k*-mers only; (3) and the histogram onto a parallel iteration. It reads the reads in parallel, in 16 MB chunks. Its table has 2^27^ buckets (60.1 GB), sized from `PanGenie`’s default estimate of 3 × 10^9^ *k*-mers. The graph’s 3.58 billion *k*-mers fill 83% of its slots, as they do `Jellyfish`’s. `PanGenie`’s own code builds with its own flags (-O3, with assertions on) in every variant, and *k*ache-hash’s library with its own. Built with -O3 `-march=native -DNDEBUG`, the release ran only 0.5% faster in sequencer order (Section D.2), so its flags do not hold it back. The control’s counter is `Jellyfish`’s table, configured as `PanGenie` configures it, behind the same parallel reader and histogram scan.

#### Runs

The runs use `PanGenie`’s own 1kGP command, `PanGenie -f <index> -i <reads> -s HG00096 -t 24 -j 24`, pinned to 24 cores, 12 per socket, with the inputs in page cache. All 54 runs, in both read orders, through both pipes and at every thread count, wrote byte-identical VCFs (but for their date line) and histograms.

#### Time and memory

Of the counting time saved over the release’s timed build, *k*ache-hash’s table saves the control’s time minus its own (Table 8). The parallel scan saves the release’s histogram time minus the control’s, and the parallel reader saves the rest. Building the panel’s index, which `PanGenie` does once per panel, peaked at 106.2 GB.

**Table 8.**
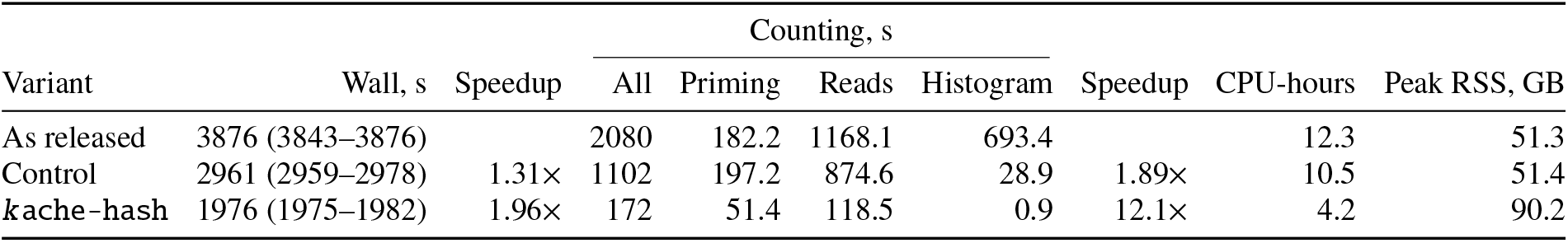
PanGenie genotyping HG00096 with the reads as distributed, by variant. Wall time is the median of three runs, with their minimum and maximum, and the speedups are over the release. CPU time is user and system time. Counting is `PanGenie`’s counting stage, and its phases are the counters’ own timers (the release’s, from a build that times them, one run).

|  |  |  | Counting, s |  |  |  |  |  |  |
|---|---|---|---|---|---|---|---|---|---|
| Variant | Wall, s | Speedup | All | Priming | Reads | Histogram | Speedup | CPU-hours | Peak RSS, GB |
|---|---|---|---|---|---|---|---|---|---|
| As released | 3876 (3843–3876) |  | 2080 | 182.2 | 1168.1 | 693.4 |  | 12.3 | 51.3 |
| Control | 2961 (2959–2978) | 1.31× | 1102 | 197.2 | 874.6 | 28.9 | 1.89× | 10.5 | 51.4 |
| *k*ache-hash | 1976 (1975–1982) | 1.96× | 172 | 51.4 | 118.5 | 0.9 | 12.1× | 4.2 | 90.2 |

### D.2 PanGenie: Counting Threads

Table 9’s runs take the read pairs sorted into the sequencer’s order by their Illumina names, without the locality of 1kGP’s genome-ordered files. At 24 threads, this order cuts *k*ache-hash’s counting speedup from 12.1× to 9.49×, but the whole run’s only from 1.96× to 1.95×. With 8 counting threads, `PanGenie`with *k*ache-hash genotypes the sample 1.50× faster than the release with 32. The release’s counting levels off at its single-threaded histogram scan.

**Table 9.**
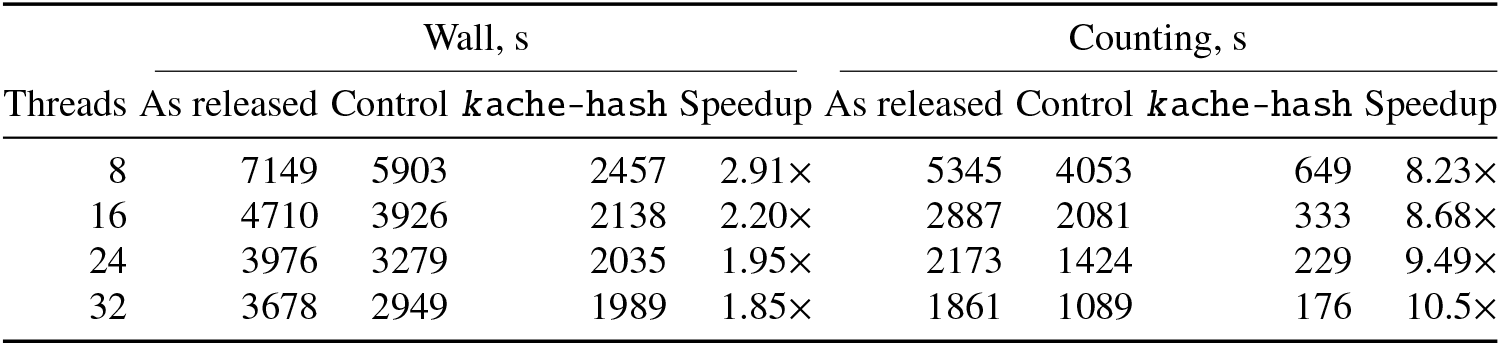
PanGenie genotyping HG00096 with the reads in sequencer order, by the counting threads (-j), one run each; the other stages keep 24 threads, and at 32 threads the runs take all 32 cores. The 24-thread line is the median of three runs, and the release’s 8-thread line the mean of two. It likely overstates the release’s time by about 8%, as part of `Jellyfish`’s table got 4 KB pages. The speedups are *k*ache-hash’s over the release.

### D.3 PanGenie: Reads Through a Pipe

`PanGenie`’s 1kGP genotyping streams each sample’s reads from gzip through `gunzip` [11]. We ran the release and *k* ache-hash this way once each, with the reads as distributed, and *k*ache-hash once more through `pigz -dc -p 8`, on the 8 cores that `PanGenie` leaves free. The release runs as fast through `gunzip` as from the file (3857 s). On its own, `gunzip` takes 708 s, four times *k*ache-hash’s counting, so the pipe bounds *k*ache-hash. It runs in 2644 s through `gunzip`, 1.46× faster than the release, and in 2144 s through `pigz`, 1.80× faster.

### D.4 K2Rmini: Integration and Runs

#### Integration

The fork [18] adds *k*ache-hash to `K2Rmini` 1.0.0 behind a new option, `--index kache`, and leaves the filter as released. Every thread builds the index. The threads take turns on items of about 4 Mbp of the memory-mapped pattern file, which overlap by k − 1 bases. The minimizers go into 1,024 shards of 1.0.0’s minimizer set. A read’s *k*-mer is a hit only if its mask in *k*ache-hash has the bit of the *k*-mer’s orientation in the read. *k*ache-hash is v0.7.0, through its C interface and its Rust bindings, under Rust 1.98.1 and GCC 11.5.

#### The fixed control

The fixed control changes only the hash of 1.0.0’s minimizer set, to Murmur3’s 64-bit finalizer in place of FxHash (rustc-hash 2.1.3). FxHash spreads keys well over up to 2^26^ buckets, as its documentation notes. Beyond that, the top bits of a bucket’s index depend only on a minimizer’s first bases. Against the human genome with its reverse complement, the set grows to 2^30^ and then 2^31^ buckets, for 764 million minimizers. Its inserts then take 1834.7 s in the release, against 77.6 s in the fixed control (Table 11). The *E. coli* genomes’ and the benchmark’s minimizer sets stay within 2^26^ buckets until the final reserve, or hold random bases. On them, the fixed control runs as the release does: in 1854 s against 1843 s on the gut reads, and 0.1–3.4% slower on the benchmark’s largest sets, in one run each with 8 threads. Of the index time that *k*ache-hash saves over the fixed control, its *k*-mer set accounts for 77% against the human genome and 72% against the *E. coli* genomes (Table 11).

#### Exactness

On every task, *k*ache-hash keeps a subset of the release’s reads. The fixed control keeps exactly the release’s (Table 10). An exact reference stores the *k*-mers’ 2-bit codes. On the benchmark’s positive sets of 2^20^, 2^22^, 2^24^ and 2^26^ patterns, it keeps exactly *k*ache-hash’s reads, with the same counts. The release keeps 1, 0, 3 and 48 reads more.

**Table 10.**
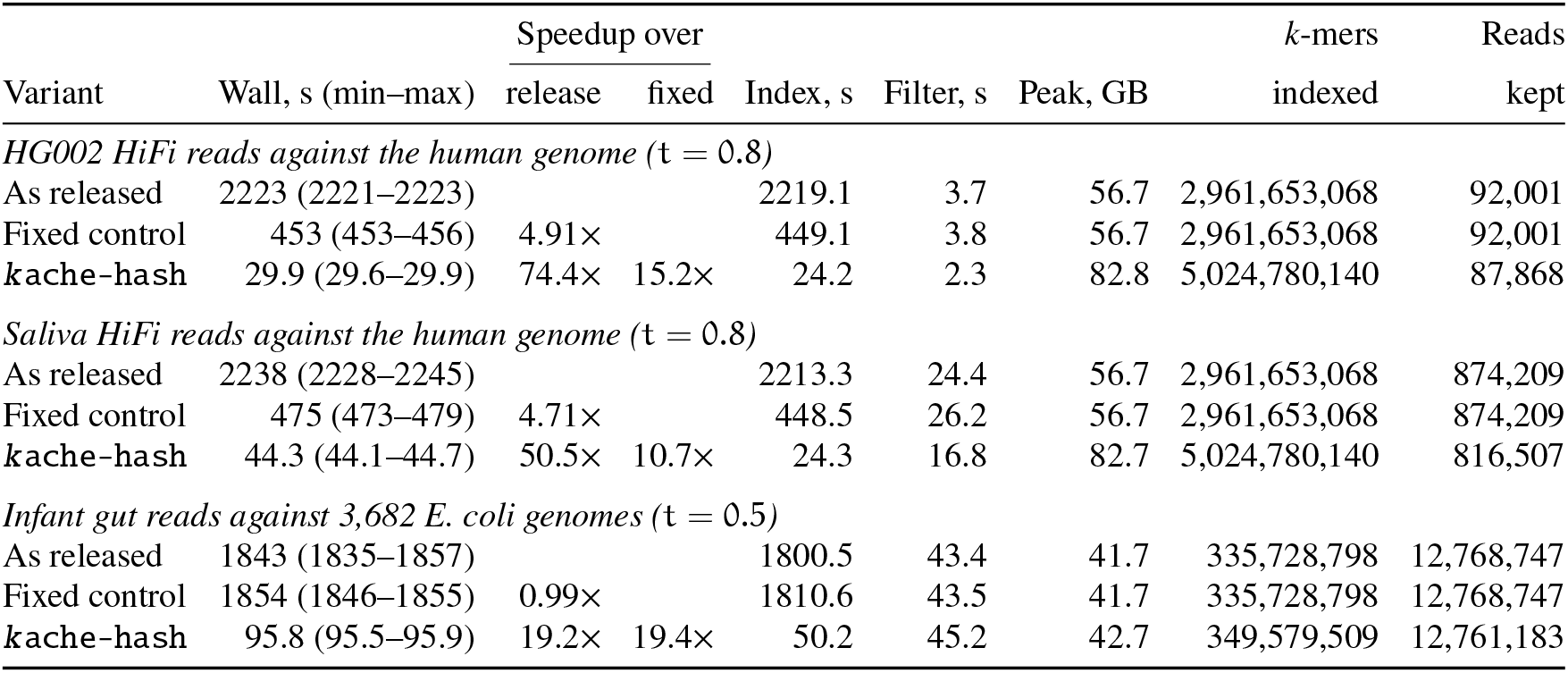
K2Rmini on the three tasks with 32 threads: as released, as the fixed control, and with *k*ache-hash. Wall time is the median of three runs, with their minimum and maximum. The index and filter times are `K2Rmini`’s own timers. The *k*-mers indexed are the release’s distinct 32-bit hashes, and *k*ache-hash’s exact *k*-mers in their forward orientations. The reads kept are from the identity runs.

| Variant | Wall, s (min–max) | Speedup over |  | Index, s | Filter, s | Peak, GB | *k*-mers | Reads |
|---|---|---|---|---|---|---|---|---|
|  |  | release | fixed |  |  |  | indexed | kept |
| *HG002 HiFi reads against the human genome (*t = 0.8*)* |  |  |  |  |  |  |  |  |
| As released | 2223 (2221–2223) |  |  | 2219.1 | 3.7 | 56.7 | 2,961,653,068 | 92,001 |
| Fixed control | 453 (453–456) | 4.91× |  | 449.1 | 3.8 | 56.7 | 2,961,653,068 | 92,001 |
| kache-hash | 29.9 (29.6–29.9) | 74.4× | 15.2× | 24.2 | 2.3 | 82.8 | 5,024,780,140 | 87,868 |
| *Saliva HiFi reads against the human genome (*t = 0.8*)* |  |  |  |  |  |  |  |  |
| As released | 2238 (2228–2245) |  |  | 2213.3 | 24.4 | 56.7 | 2,961,653,068 | 874,209 |
| Fixed control | 475 (473–479) | 4.71× |  | 448.5 | 26.2 | 56.7 | 2,961,653,068 | 874,209 |
| kache-hash | 44.3 (44.1–44.7) | 50.5× | 10.7× | 24.3 | 16.8 | 82.7 | 5,024,780,140 | 816,507 |
| *Infant gut reads against 3,682 E. coli genomes (*t = 0.5*)* |  |  |  |  |  |  |  |  |
| As released | 1843 (1835–1857) |  |  | 1800.5 | 43.4 | 41.7 | 335,728,798 | 12,768,747 |
| Fixed control | 1854 (1846–1855) | 0.99× |  | 1810.6 | 43.5 | 41.7 | 335,728,798 | 12,768,747 |
| kache-hash | 95.8 (95.5–95.9) | 19.2× | 19.4× | 50.2 | 45.2 | 42.7 | 349,579,509 | 12,761,183 |

**Table 11.** The index’s time in seconds, by its parts. The release’s and the fixed control’s parts come from split builds that skip the inserts into the *k*-mer set, or into both sets (one run each). *k*ache-hash’s parts are its threads’ average time in each set, as parsing and hashing run inside its threads. Each whole index is the median of three runs.

| Variant | Parsing, hashing | Minimizer set | *k*-mer set | Index |
|---|---|---|---|---|
| *Against the human genome* |  |  |  |  |
| As released | 26.2 | 1834.7 | 358.2 | 2219.1 |
| Fixed control | 26.2 | 77.6 | 345.2 | 449.1 |
| kache-hash |  | 4.8 | 17.5 | 24.2 |
| *Against the E. coli genomes* |  |  |  |  |
| As released | 153.0 | 320.3 | 1327.1 | 1800.5 |
| Fixed control | 153.0 | 348.4 | 1309.2 | 1810.6 |
| kache-hash |  | 13.9 | 34.6 | 50.2 |

#### The benchmark

Up to 2^22^ patterns, `K2Rmini`’s single parser thread bounds the release’s runs. At these sizes, *k*ache-hash’s start-up costs up to 0.15 s: its threads, the minimizer set’s 1,024 shards and its table.

#### Runs

The runs filter the reads with `K2Rmini`’s command, `K2Rmini -k 31 -m 21 -t <t> -T <threads> -p <patterns> <reads> -o /dev/null`, on the server of Section 5. They are pinned to cores 0 to T − 1, with their inputs in page cache and transparent huge pages always on. Each configuration ran three times, interleaved, and each split build once. The gut reads leave out the run’s 92 reads that hold no 31 bases between runs of N. No threshold keeps such a read. One that opens a batch crashes the release.

## Footnotes

3 Keys are canonical *k*-mers (§2.2) and the complement of zero, the all-ones word, is never a canonical *k*-mer.

4 On Linux, `mremap` moves the main table’s pages to a range twice as large, without copying them.

5 `IcebergHT’`s faulty resized builds aside (§4.1).

## Notes

### Competing Interest Statement

RP is a co-founder of Ocean Genomics Inc.

### Summary of Updates

The design of the proposed data structure, along with its implementation, has been improved substantially. The results have therefore improved. Applications for the method have also been demonstrated.

### Funding

- National Institutes of Health: R01HG009937
- U.S. National Science Foundation: OAC 2517201, OAC 2513656
- Chan Zuckerberg Initiative (United States): 2022-311195, 2024-34282

